# Sex-biased admixture and assortative mating shape genetic variation and influence demographic inference in admixed Cabo Verdeans

**DOI:** 10.1101/2020.12.14.422766

**Authors:** Katharine L Korunes, Giordano Bruno Soares-Souza, Katherine Bobrek, Hua Tang, Isabel Inês Araújo, Amy Goldberg, Sandra Beleza

## Abstract

Genetic data can provide insights into population history, but first we must understand the patterns that complex histories leave in genomes. Here, we consider the admixed human population of Cabo Verde to understand the patterns of genetic variation left by social and demographic processes. First settled in the late 1400s, Cabo Verdeans are admixed descendants of Portuguese colonizers and enslaved West African people. We consider Cabo Verde’s well-studied historical record alongside genome-wide SNP data from 563 individuals from 4 regions within the archipelago. We use genetic ancestry to test for patterns of nonrandom mating and sex-specific gene flow, and we examine the consequences of these processes for common demographic inference methods and for genetic patterns. Notably, multiple population genetic tools that assume random mating underestimate the timing of admixture, but incorporating non-random mating produces estimates more consistent with historical records. We consider how admixture interrupts common summaries of genomic variation such as runs-of-homozygosity (ROH). While summaries of ROH may be difficult to interpret in admixed populations, differentiating ROH by length class shows that ROH reflect historical differences between the islands in their contributions from the source populations and post-admixture population dynamics. Finally, we find higher African ancestry on the X chromosome than on the autosomes, consistent with an excess of European males and African females contributing to the gene pool. Considering these genomic insights into population history in the context of Cabo Verde’s historical record, we can identify how assumptions in genetic models impact inference of population history more broadly.

## Introduction

Patterns of genetic variation are often used to infer human population histories, such as fluctuations in population size, the timing of gene flow, and changes in population structure (Loh et al. 2013; Schiffels and Durbin 2014; Browning and Browning 2015; Goldberg and Rosenberg 2015; Zaitlen et al. 2017; Browning et al. 2018; Ni et al. 2019). These studies have been informative about ancient and modern populations that may lack historical records (Skoglund et al. 2012; Lazaridis et al. 2014; Busby et al. 2015; Fernandes et al. 2019; Font-Porterias et al. 2019). However, by necessity, population-genetic models make a variety of simplifying assumptions that often neglect the demographic and social dynamics of human populations. For example, many classical models assume random mating, but within empirical populations mating is often nonrandom, known as assortative mating. Positive assortative mating in human populations can keep haplotype blocks together, affecting evolutionary processes and our ability to infer population history (Risch et al. 2009; James Y Zou et al. 2015; Sebro et al. 2017). Further, male and female demographic processes can differ and vary over time. Comparisons between the X chromosome and autosomes have shown that population sizes and migration rates differ between males and females of many human populations (Wilkins and Marlowe 2006; Ramachandran et al. 2008; Bustamante and Ramachandran 2009; Keinan et al. 2009; Arbiza et al. 2014).

The challenges of inferring human demographic history are particularly apparent on short timescales of tens of generations, where changes in allele frequencies may be difficult to observe and biased by dynamics not captured by classic models. Admixed populations provide an opportunity to examine evolutionary processes on short timescales using admixture linkage-disequilibrium (LD) structure rather than potentially small changes in allele frequencies. Understanding the patterns generating the distribution of ancestry within an admixed population is also important as genomic data has shown how widespread admixture is throughout human evolution (Hellenthal et al. 2014; Lazaridis et al. 2014; Ruiz-Linares et al. 2014; Busby et al. 2015; Triska et al. 2015; Busby et al. 2016; Laso-Jadart et al. 2017; Korunes and Goldberg 2021).

In admixed populations, nonrandom mating can be driven by factors that correlate with genetic ancestry. For example, spouse pairs within multiple Latino populations are correlated in their genomic ancestry (Risch et al. 2009; James Y Zou et al. 2015). When mating patterns lead to a positive correlation in genomic ancestry, this specific type of nonrandom mating, referred to as ancestry-assortative mating, can increase the likelihood that individuals in mating pairs share recent common ancestors. Recent work has begun to assess the implications of these patterns, including effects on common demographic inference strategies and on genetic/phenotypic variation (Ruiz-Linares et al. 2014; Sebro et al. 2017; Zaitlen et al. 2017). One genetic consequence of positive ancestry-assortative mating is an increase in long runs of homozygosity (ROH), which arise when long tracts of homozygous genotypes occur due to inheritance of identical by descent (IBD) haplotypes from recent common ancestors. Across modern human populations, ROH are common and typically range in length from kilobases to megabases (McQuillan et al. 2008; Kirin et al. 2010; Pemberton et al. 2012; Blant et al. 2017). In the field of population genetics, we are just beginning to learn about ROH dynamics and expectations in admixed populations, which do not necessarily fit expectations built from homogenous populations (Pemberton et al. 2012; Mooney et al. 2018). For example, we know that distinct length classes of ROH reflect demographic factors from different timescales, but as we explore here, it is unclear what these distributions of ROH mean in populations that have undergone major demographic transitions such as recent admixture. Patterns of genetic variation can also be shaped by differences in male and female demographic processes. In many human populations, population sizes and migration rates differ between males and females, leading to differences in ancestry patterns on the X chromosome vs the autosomes (Wilkins and Marlowe 2006; Ramachandran et al. 2008; Bustamante and Ramachandran 2009; Keinan et al. 2009; Arbiza et al. 2014).

Here, we use the admixed population of Cabo Verde as a case study to investigate the patterns of genetic variation left by multifaceted social and demographic processes in humans. Admixed populations of African ancestry, such as the population of Cabo Verde, are often excluded in the context of medical genetics and human evolutionary genetics, despite their importance and widespread global presence (Popejoy and Fullerton 2016; Landry et al. 2018; Popejoy et al. 2018). The record of Cabo Verde population history is vast, including royal charters, letters from Crown officials, captains and other sea explorers’ journals, records of the economic activities in Cabo Verde (e.g., landowners’ rents and number of enslaved individuals traded), detailed ship logs of trade activity, and church censuses from the 18^th^ century on, which allows us to generate realistic assumptions about colonization and demography (Barbosa, Gilda 1997; Russell-Wood 1998; Baleno 2001; Cabral 2001; Correia e Silva 2001; Correia e Silva 2002; Cabral 2012). We consider patterns of genetic variation alongside historical records that document many aspects of Cabo Verdean history, including settlement patterns, timing, and sociocultural dynamics. Specifically, we use distributions of genetic ancestry to test for ancestry-assortative mating and sex-specific gene flow. We examine the consequences of these processes for genetic variation, such as patterns of homozygosity, and for common demographic inference methods. In turn, by elucidating the patterns of genetic variation social processes leave in human genomes for a population with a well-studied historical and ethnographic record, we can better use population-genomic methods to explore the history of populations without extensive records.

### Cabo Verdean history

The recently admixed human population of Cabo Verde presents a particularly relevant opportunity to examine how sex-biased admixture and nonrandom mating shape genetic patterns and influence demographic inference. Cabo Verde is an archipelago off the coast of Senegal, inhabited today by admixed descendants of Portuguese colonizers and enslaved West African people who settled the unpopulated islands beginning in the mid-1400s (Fernandes et al. 2003; Beleza et al. 2012; Beleza et al. 2013; Verdu et al. 2017; Laurent et al. 2022). Since the archipelago was unoccupied prior to the 1460s, we have historical knowledge of the start of admixture and the contributing source populations, with written records clearly documenting arrival times by island. Additionally, the island geography imposes population structure within Cabo Verde, and historical records document differences in population sizes, mating patterns, and social customs by island population (see Methods for historical data sources).

The settlement of the islands was influenced by island geography and ecology, and is often divided into three different temporal stages, which are associated with changes in the economy (Correia e Silva 2002). The initial settlement stage (beginning in the 1460s) corresponds to the peopling of the Southern island Santiago, followed by the nearby island of Fogo 20 years later (Fig 1). During this time, the economy was centered around large cotton plantations and expanding trade with the Senegambian coast. This trade included the exportation of great numbers of enslaved Africans to Cabo Verde, the majority of which became part of the commercial trade to the Americas (Russell-Wood 1998; Baleno 2001). In the 17^th^ century, following the decline of the slavery-based economy of Santiago and Fogo, many free farmers seeking better farming conditions migrated to the northwestern islands of Santo Antão and São Nicolau. These migrations comprised the second settlement stage. In contrast to Santiago and Fogo, these northwestern islands did not become populous economic centers (Correia e Silva 2002). During the third and final settlement stage, São Vicente in the northwestern group of islands was colonized by settlers from the neighboring islands in the mid 19^th^ century. With the opening of Mindelo’s harbor at that time, São Vicente prospered from the advent of commercial Atlantic shipping, becoming the second most important island of the archipelago after Santiago in terms of population size. In the 19^th^ century, the eastern islands Sal, Boa Vista, and Maio were also open to English and North American ships, but these islands never became densely populated (Russell-Wood 1998; Correia e Silva 2002).

**Fig 1.**
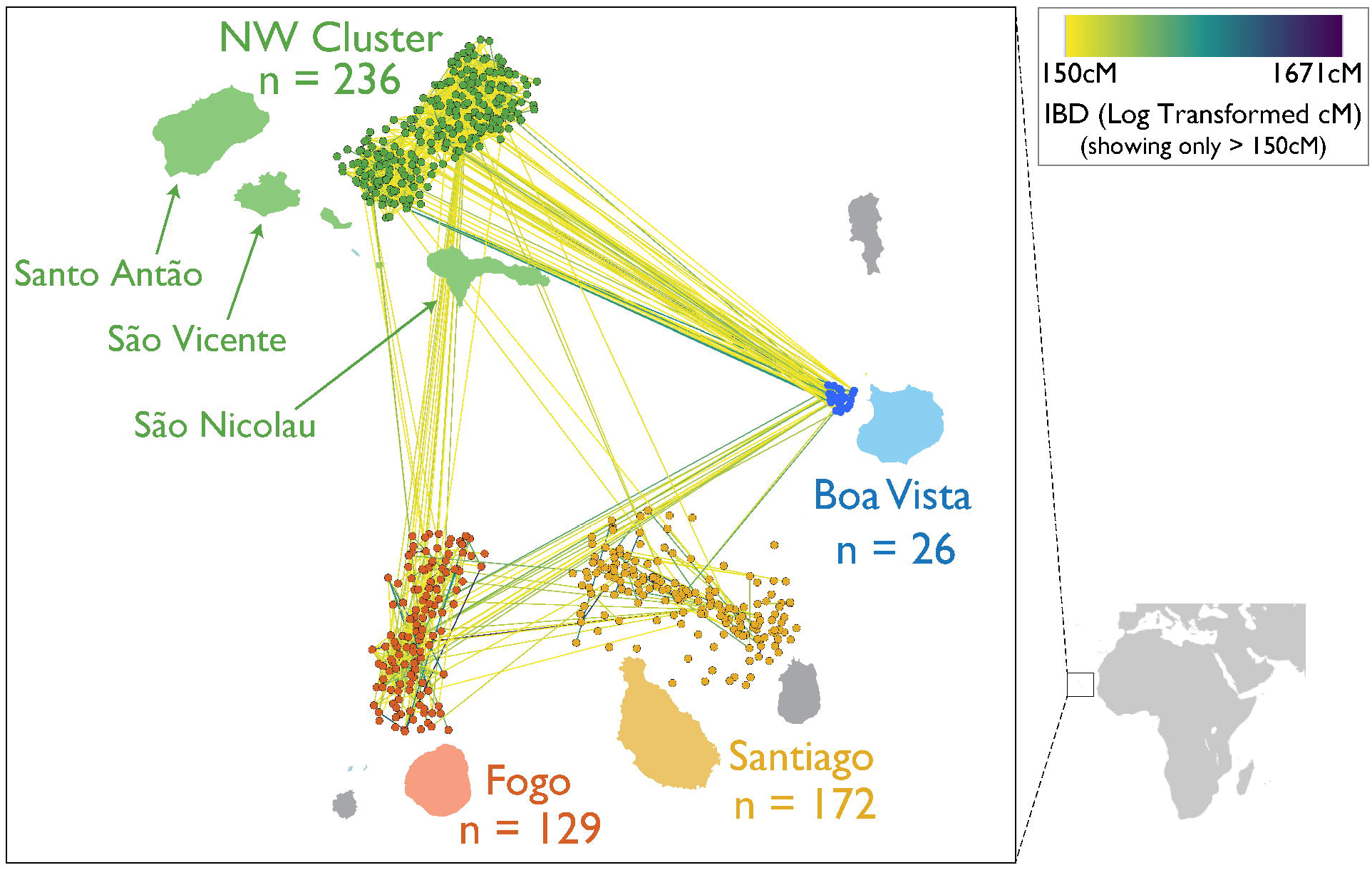
Shared ancestry between and within the islands in the context of geography. Each island has a corresponding cluster of nodes representing all sampled individuals, with the individuals localized to be adjacent to the island where they were sampled. Node placement within islands is determined by a force-directed algorithm using pairwise shared IBD, meaning that the spread of each cluster reflects the level of relatedness in each population. Edges between the nodes represent total IBD tract length for pairs of individuals sharing more than 150 cM of total IBD, with edges colored using the log transformed total IBD length.

Previous investigations of genetic ancestry in Cabo Verde (Beleza et al. 2012; Beleza et al. 2013; Verdu et al. 2017) have shown extensive admixture in the archipelago and differences in the mean African genetic ancestry across islands, which may reflect the differences in the settlement history and suggest restricted gene flow between islands. These past studies of Cabo Verdean genetic admixture have explored how genetic ancestry connects to phenotypic variation such as skin and eye color (Beleza et al. 2012; Beleza et al. 2013), and to cultural information such as linguistic variation (Verdu et al. 2017). Some earlier analyses of post-admixture population structure were based on Y-chromosome diversity (Beleza et al. 2012), giving insight into male-specific demographic processes. Here, we examine how historical and social processes influence genetic variation and population-genetic inference of demography. We consider how the distributions of segments of DNA shared identically by descent within and among individuals (as measured by IBD, kinship, and ROH) relate to patterns of genetic ancestry in an admixed population. We integrate several population genomic approaches using autosomal and X-chromosome ancestry and genetic variation patterns to infer the sex-specific demographic history of the last ∼20 generations since founding, and we consider the potential biases of ancestry-assortative mating on the inference of admixture timing. Overall, we provide insights into the population history of Cabo Verde and demonstrate the need to better understand how demographic inference methods and common summaries of genomic variation operate in admixed populations.

## Methods

### Study population and ancestry reference panels

We used genotype data from Beleza et al. (2013) (Beleza et al. 2013), which included 563 Cabo Verdeans from the following regions of the archipelago: Santiago (n = 172), Fogo (n = 129), Boa Vista (n = 26), and the three northwestern islands in aggregate (Northwest Cluster; n = 236) (Figure 1; Supp Fig 1). The genotype data exclude cryptic related individuals (first-degree relatives) identified by kinship analyses at the time of sampling. In accordance with historical records and previous work showing that Iberian and Senegambian populations are suitable proxies for the ancestry sources of Cabo Verde (Verdu et al. 2017), we leveraged data from the 1000 Genomes Project to estimate admixture proportions and call local ancestry in the Cabo Verde individuals, as described below. We merged the Cabo Verde genotypes with genotypes from 107 GWD (Gambian in Western Division - Mandinka) and 107 IBS (Iberian Population in Spain) samples called from high-coverage resequencing data released through the International Genome Sample Resource (Clarke et al. 2017; Fairley et al. 2020). Our final merged dataset consisted of 884,656 autosomal and 20,967 X chromosome SNPs shared between the Cabo Verde samples and the reference samples, with average missingness rates of 0.0017 by SNP for autosomes, and 0.0024 for the X chromosome.

### Historical records

Throughout our analyses, we draw comparisons between genetic results and historical records. We used primary historical documents, mainly historical letters to the Portuguese Crown, documenting demographic characteristics of the islands across time. Our analyses of the historical data took into consideration the generalized interpretations of historian scholars such as Correia e Silva (2001; 2002) (Correia e Silva 2001; Correia e Silva 2002), Cabral (2001; 2012) (Cabral 2001; Cabral 2012) and Baleno (2001) (Baleno 2001). These sources document the dates of the stages of settlement mentioned above, along with associated with changes in the economy. We refer to the interpretation of genealogical data by Cabral (2012) (Cabral 2012) and Barbosa (1997) (Barbosa, Gilda 1997) for evidence of the structure of the Fogo’s society and for evidence of first-cousin marriages within Fogo (Barbosa, Gilda 1997).

### Characterization of ancestry

To produce estimates of admixture proportions, we first performed unsupervised clustering of the samples using ADMIXTURE v1.3.0 (Alexander et al. 2009) (see Supp Table 1 for a summary of all computational methods used in this study). ADMIXTURE was run separately for the autosomal and X chromosome datasets after pruning based on linkage disequilibrium (LD) using the indep-pairwise option of PLINK v1.9 with a 50-SNP sliding window incremented by 10 SNPs, and an LD threshold of r2 = 0.5 (Purcell et al. 2007). This pruned dataset included 514,551 autosomal and 12,706 X chromosome SNPs. We estimated individual ancestries by averaging over ten independent unsupervised ADMIXTURE runs using K = 2, given the historical and genetic support for two source populations (Beleza et al. 2013; Verdu et al. 2017). Using the LD-pruned dataset, we also visualized the data using PLINK’s principal component analysis (--pca) function, confirming that the samples cluster approximately based on their geographic memberships, and that West African vs European ancestry clearly separates on the first principal component (Supp Fig 1). We phased all samples using SHAPEIT2 (Delaneau et al. 2013). After running SHAPEIT -check to exclude sites not contained within the reference map, we ran SHAPEIT to yield phased genotypes at 881,279 autosomal SNPs and 20,793 X chromosome SNPs. We then called local ancestry with RFMix v1.5.4 (Maples et al. 2013) PopPhased under a two-way admixture model using the West African and European reference genotypes described above. These local ancestry calls are publicly available via Zenodo (Hamid et al. 2020). Ancestry proportions estimated with ADMIXTURE and RFMix here are highly correlated with results from Beleza et al. (2013) (Beleza et al. 2013), using the same genotype data but with different software (*frappe* (Tang et al. 2005) and SABER (Tang et al. 2006)) and an older reference dataset (Supp Fig 2). To test whether phasing and local ancestry calls were robust against the choice of reference panels, we repeated the SHAPEIT and RFMix steps using a reference dataset composed by all African (ACB, ASW, ESN, GWD, LWK, MSL, and YRI) and European (CEU, FIN, GBR, IBS, and TSI) populations available from the high-coverage resequencing data released by the 1000 Genomes Project (Clarke et al. 2017; Fairley et al. 2020). The resulting local ancestry calls with all AFR and EUR reference panels correlated closely with calls using GWD and IBS as reference panels (Supp Fig 2C). We also performed local ancestry calling using ELAI, a method that performs both phasing and local ancestry assignment (Guan 2014). Again using the IBS and GWD reference genotypes described above, we ran ELAI under a two-way admixture model using the following parameters: -mg (number of generations) 20, -s (EM steps) 30, -C (upper clusters) 2, and -c (lower clusters) 10. The resulting local ancestry calls from ELAI correlated closely with calls from RFMix (Supp Fig 2D).

### Inference of admixture timing

We applied three distinct strategies for estimating the timing of the onset of admixture in Cabo Verde: ALDER (Loh et al. 2013), MultiWaver 2.0 (Ni et al. 2019), and a method based on patterns of linkage disequilibrium between local ancestry tracts (Zaitlen et al. 2017). These methods differ in their strategies for inferring admixture timing and subsequently differ in their input data and in their underlying assumptions. Specifically, ALDER uses the extent of LD decay among neighboring loci to infer mixture proportions and dates; MultiWaver uses the length distribution of ancestral tracks; and the strategy of Zaitlen et al. (2017) uses patterns of linkage disequilibrium between local ancestry tracts as described below. To prepare the input data for each of these methods, we first converted the genotypes to EIGENSTRAT format using *convert* (Patterson et al. 2006; Price et al. 2006). We then used ALDER to date admixture timing on each island using default parameters with mindis = 0.005 and source populations GWD and IBS. We ran MultiWaver using the ancestry tracts inferred by RFMix, default parameters, and 100 bootstraps. Finally, we followed the pipeline of Zaitlen et al. (2017) (Zaitlen et al. 2017) and its accompanying scripts to use the RFMix local ancestry calls to measure local ancestry disequilibrium (LAD) decay in 10 Mb windows, overlapping by 1 Mb. Specifically, we started with the first SNP on each autosome, used the 10 Mb window end point to identify the SNP closest to the inside of this boundary, and then used local ancestry calls at these positions to determine LAD. We repeated this process along each chromosome to obtain LAD in 279 autosomal windows. Using possible values of admixture generations ranging from 5-25, we determined the best fit using island-specific mean LAD decay over the 279 autosomal windows, assortative mating parameters estimated with ANCESTOR (see below), a starting autosomal admixture proportion of 0.65, and either no migration or with migration (migration rate = 0.01).

### Tests for ancestry-assortative mating

We tested for ancestry-assortative mating over the last generation using ANCESTOR (James Y. Zou et al. 2015; James Y Zou et al. 2015). ANCESTOR uses phased local ancestry tracts in the current generation to estimate the ancestral proportions of the two parents of each individual. Specifically, ANCESTOR uses a framework of pooled semi-Markov processes to infer the parameters of the parental ancestries for individuals in an admixed population. Thus, we provided the local ancestry tracts inferred using RFMix as input to ANCESTOR’s ancestor.py script. We used the inferred parental ancestries to test for assortative mating seen as a positive correlation in ancestry between inferred mating pairs (Parent 1 vs Parent 2 African ancestry proportion), and we used the strength of the observed positive correlation (Pearson’s R for each population, using the median value from all chromosomes) as the assortative mating parameter in our application of the LAD-based method for inferring admixture timing. We also examined evidence of nonrandom mating by estimating genetic relatedness within Cabo Verde using both IBD (see below) and kinship coefficients (Ochoa and Storey 2019 May).

### Identification of IBD tracts and ROH

Following the analysis pipeline of S. R. Browning et al. (2018) (Browning et al. 2018), we inferred segments of IBD using the haplotype-based IBD detection method Refined IBD (Browning and Browning 2013), and we estimated ancestry-specific population size using IBDNe (Browning and Browning 2015). After running Refined IBD, we used its gap-filling utility to remove gaps between segments less than 0.6 cM that had at most one discordant homozygote. We then filtered out IBD segments smaller than 5 cM, as short segments below this threshold are difficult to accurately detect (Nelson et al. 2020). To visualize IBD sharing in Fig 1 and Supp Fig 3, the sum of IBD shared in each pairwise comparison of individuals was plotted by using Cytoscape 3.8 to group nodes by island, position nodes within each island according to a prefuse force-directed algorithm, and scale the color of edges based on log-transformed total IBD length (Shannon et al. 2003).

We called ROH using GARLIC v.1.1.6 (Szpiech et al. 2017), which implements the ROH calling pipeline of Pemberton et al. (2012) (Pemberton et al. 2012). We performed this analysis separately for the autosomes and X chromosome. Based on the pipeline described in Szpiech et al. (2017), we used a single constant genotyping error rate of 0.001, we allowed GARLIC to automatically choose a window size for each population (--auto-winsize), and we used the resample flag to mitigate biases in allele frequency estimates caused by differing sample sizes. This resulted in GARLIC selecting a window size of 50 SNPs in each of the Cabo Verde regions and in GWD, and a window size of 40 for IBS. Using a three-component Gaussian mixture model, GARLIC classified ROH into three length groups: small/class A, medium/class B, and long/class C ROH. Across all populations, class A/B and class B/C size boundaries were inferred as approximately 300 kb and 1 Mb, respectively (see Supp Table 2 for population-specific parameters, including LOD cutoffs and size boundaries). Using only the females, we then classified ROH for the X chromosome.

Since levels of ROH partitioned by size class are dependent on the size classification cutoffs, we assessed concordance of our results by repeating size classification using the --size-bounds flag in GARLIC to impose four additional sets of size class boundaries. We first classified ROH based on the commonly used rule of thumb that a time depth of *m* meioses is expected to give a mean IBD segment length of 100/*m* cM (Thompson 2013). In this case, common ancestors within the past 20 generations (post-admixture) would give an expected IBD segment length of approximately 2.5 Mb or longer, so we applied 2.5 Mb as the shorter/long ROH boundary. We also collected the population-specific length classification cut-offs from the 64 worldwide populations provided by Pemberton et al. (2012), which represents an extensive range of classification schemes. We repeated ROH-calling with GARLIC using the minimum, mean, and maximum of the 64 population-specific boundaries reported in Pemberton et al. (2012).

### Estimation of sex bias

To examine sex bias, we applied a mechanistic model of sex-biased admixture (Goldberg and Rosenberg 2015) to infer admixture parameters under a model of constant migration. We ran the script provided by Goldberg and Rosenberg to compare their model-based expectations of X and autosomal ancestry with observed ancestry in our data using the following parameters: grid increment size (0.02), the observed autosomal and X admixture proportions (based on ADMIXTURE estimates described above), the proportion of females in the sample, and the percentile cutoff for parameters to keep (0.1%). We performed this analysis for all 563 Cabo Verdean individuals pooled together, and then repeated the analysis for each island independently.

## Results

### Patterns of shared ancestry reflect the colonization history of the islands

Using genome-wide SNP data from 563 individuals, we examined four island regions of Cabo Verde (Fig 1; Supp Fig 1), which differ in their settlement histories. The partitioning of the considered island regions was also supported by genetic patterns. Namely, these regions showed quantitatively distinct distributions of IBD tracts, as we explore below, and were supported by clustering patterns in PCA (Supp Fig 1), in which the three northwestern islands grouped together (and with the eastern island Boa Vista) despite the different settlement time of São Vicente Island.

Global ancestry proportions reflect extensive European and West African admixture and concur with previous results (Beleza et al. 2013) (Supp Fig 2), with the median West African autosomal ancestry in Cabo Verdeans varying by island from 50% in Fogo, 56% in the northwestern islands in aggregate (the Northwest Cluster), 63% in Boa Vista, and 75% in Santiago. These inter-island differences alongside historical knowledge provide a comparative dataset for understanding how ancestry patterns can be used to infer demographic processes. Further, the known bound on admixture timing and the relative simplicity of the admixture history on the islands allows genetic ancestry to be decomposed into two source populations, which makes Cabo Verde a case study closer to population genetic models of admixture than most worldwide human populations.

The process of admixture can influence measures such as IBD and ROH that are often used to inform inference of population history, and theoretical expectations for these measures are less clear for admixed populations compared to homogeneous populations. Thus, we use multiple methods to examine relatedness in Cabo Verde, and we use this case study to underscore the need for further empirical and theoretical work to understand the dynamics of IBD and ROH in admixed populations. Patterns of IBD within and between populations provide opportunities to examine common ancestry based on the number and sizes of segments of IBD. Using these summaries of IBD to examine relatedness between and within regions of Cabo Verde, we found that patterns of shared ancestry reflect the successive settlement history of the islands. Notably, Santiago has the lowest mean number and total length of IBD segments between individuals (Supp Fig 4-5). In contrast, we observe the highest levels of IBD within and between the Northwest Cluster and Boa Vista.

To summarize patterns of IBD within and between the four island regions, we built a network of relationships with pairs of individuals connected if they have IBD levels in the top 3% of the Cabo Verde IBD distribution (Fig 1; see Supp Fig 3 for lower levels of IBD and within-island networks). In a non-admixed population, these edges roughly correspond to 4^th^-degree relatives or closer (e.g., great-great-grandparents, great-great-grandchildren, great-aunts/uncles, or grand-nieces/nephews). At this level of relatedness, we find that the presence of pairs of individuals within and between islands connected by an edge is common, suggestive of recent relatives both within and across islands. However, relatedness parameters estimated under simple models of structure likely do not hold in many human populations, such as recently admixed populations. Kinship estimated under a framework (Ochoa and Storey 2019 May) specifically designed for arbitrary population structures (Supp Fig 6A) is consistent with both the overall trends in IBD tracts (Supp Fig 4-5), and supports the presence of a subset of very closely related individuals within the data.

The lower levels of IBD (Fig 1, Supp Fig 3-5) and lower kinship estimates (Supp Fig 6) observed in Santiago are consistent with expectations based on the island’s history, as Santiago was the first Cabo Verdean island to be founded and has the largest population size. Santiago is also likely to have relatively high genetic diversity as a result of having the highest proportion of West African ancestry, based on the expectation that kinship in human populations generally reflects serial bottlenecks due to dispersal from Africa (Tishkoff and Williams 2002; Reed and Tishkoff 2006). In contrast, the high levels of IBD within and between the Northwest Cluster and Boa Vista reflect well-documented serial founding and migrations during the settlement of the islands. In general, kinship estimates also reflect the colonization history of the islands, with Cabo Verdeans of greater European ancestry sharing greater kinship compared to individuals with greater West African ancestry (Supp Fig 6). Exceptions to this trend were most noticeable in individuals from Fogo and the Northwest Cluster, where it was most apparent that some observed relatedness patterns do not result in smooth ordering of global ancestry proportions (Supp Fig 6B). This observation led us to hypothesize that other sociocultural processes beyond proportions of ancestry, such as nonrandom mating, may drive patterns of relatedness in Cabo Verde.

### Nonrandom mating shapes patterns of ancestry and influences demographic inference

To test for ancestry assortative mating within Cabo Verde, we examined whether the genomic ancestries of individuals in mating pairs correlate with each other. To this end, we applied ANCESTOR to computationally infer the parental ancestry proportions that likely preceded the ancestry haplotypes we observe today. We repeated this analysis for each chromosome independently to prevent uncertainty introduced by matching interchromosomal haplotypes. As an example, the inferred ancestries of the parental haplotypes that likely preceded chromosome 7 are significantly positively correlated for all islands except for Boa Vista (Fig 2A). We observed similar results across the full set of autosomes (Fig 2B). We found that the inferred ancestries of mating pairs in the previous generation are positively correlated, and that these correlations differ significantly from expectations under random mating (random sampling empirical distributions of parental haplotypes into pairs, shown in Supp Fig 7).

**Fig 2.**
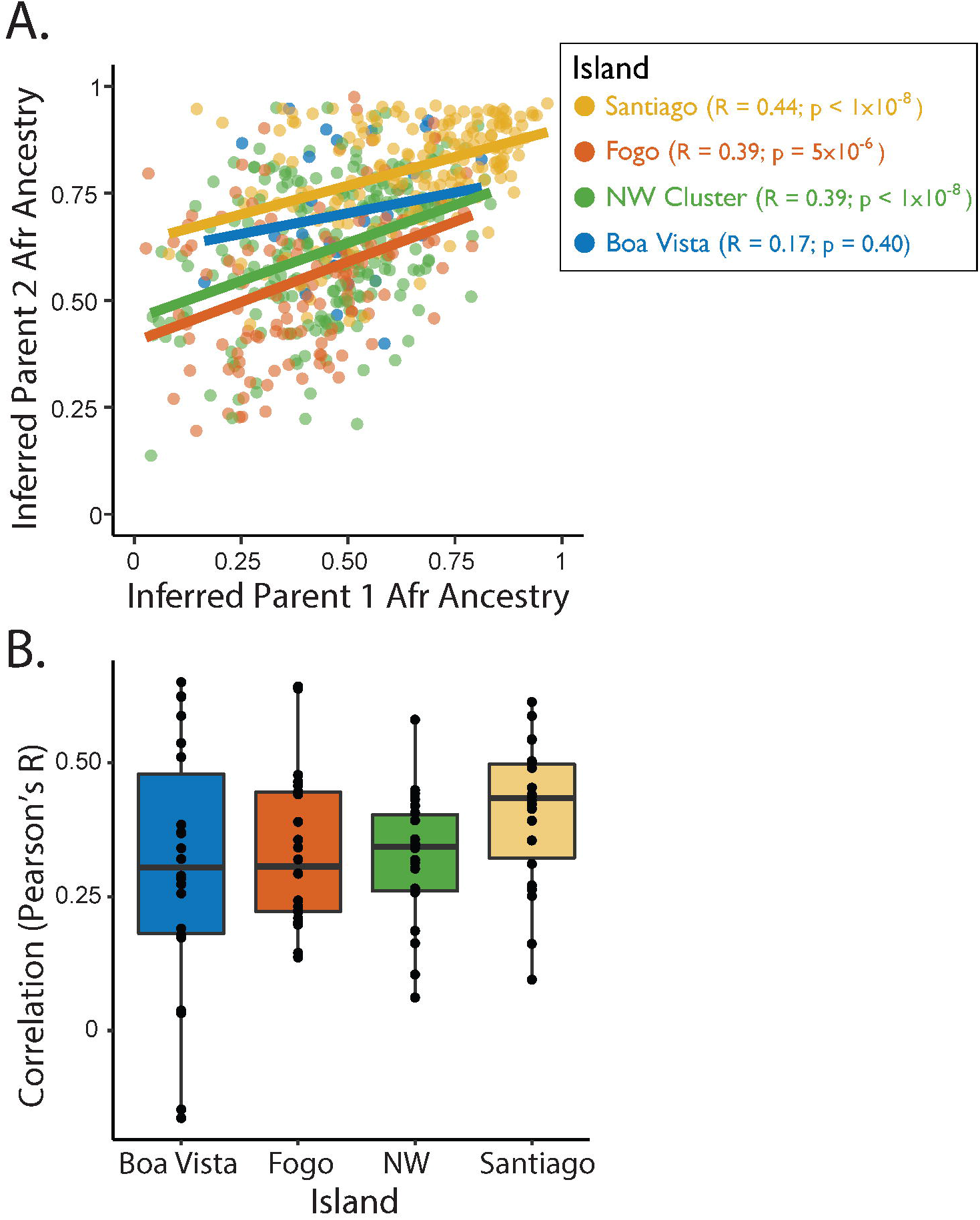
Correlation between inferred parental ancestries. (A) For each Cabo Verdean individual, the inferred Parent 1 vs Parent 2 ancestry proportions are shown for an example chromosome (Chromosome 7), colored by island. (B) shows the set of correlation coefficients between inferred parental ancestries for each autosomal chromosome.

We next considered how the observed genetic evidence of assortative mating can be leveraged in the context of inferring population history. To estimate the timing of the onset of admixture, we applied the method of Zaitlen et al. (2017) (Zaitlen et al. 2017), which describes the decay of local ancestry disequilibrium (LAD) as a function of assortative mating strength, migration rate, recombination rate, and the number of generations since admixture began. Using this LAD-based method, we inferred admixture timing under random mating and under our empirical estimates of ancestry-assortative mating strength (Fig 3). The LAD-based method produced older estimates of admixture timing under a model including both assortative mating and constant migration (Fig 3, Supp Fig 8), with increasingly older timing estimates as assortative mating strength increases (Supp Fig 8). Under the same set of migration rates (0 or 1% per generation), when we varied the strength of ancestry-assortative mating from a situation of random mating to a situation of strong ancestry-assortative mating, the models that considered substantial ancestry-assortative mating (parental correlation in ancestry > 0.3), yielded admixture timing estimates closest to the historical estimates (Fig 3, Supp Fig 8). We note that the migration rates (0 or 1% per generation) demonstrate estimates under a broad range that likely includes the historical average over time (see Methods for historical data sources), although it does not capture the fluctuations in migration rate over time that occurred within the islands (Laurent et al. 2022).

**Fig 3.**
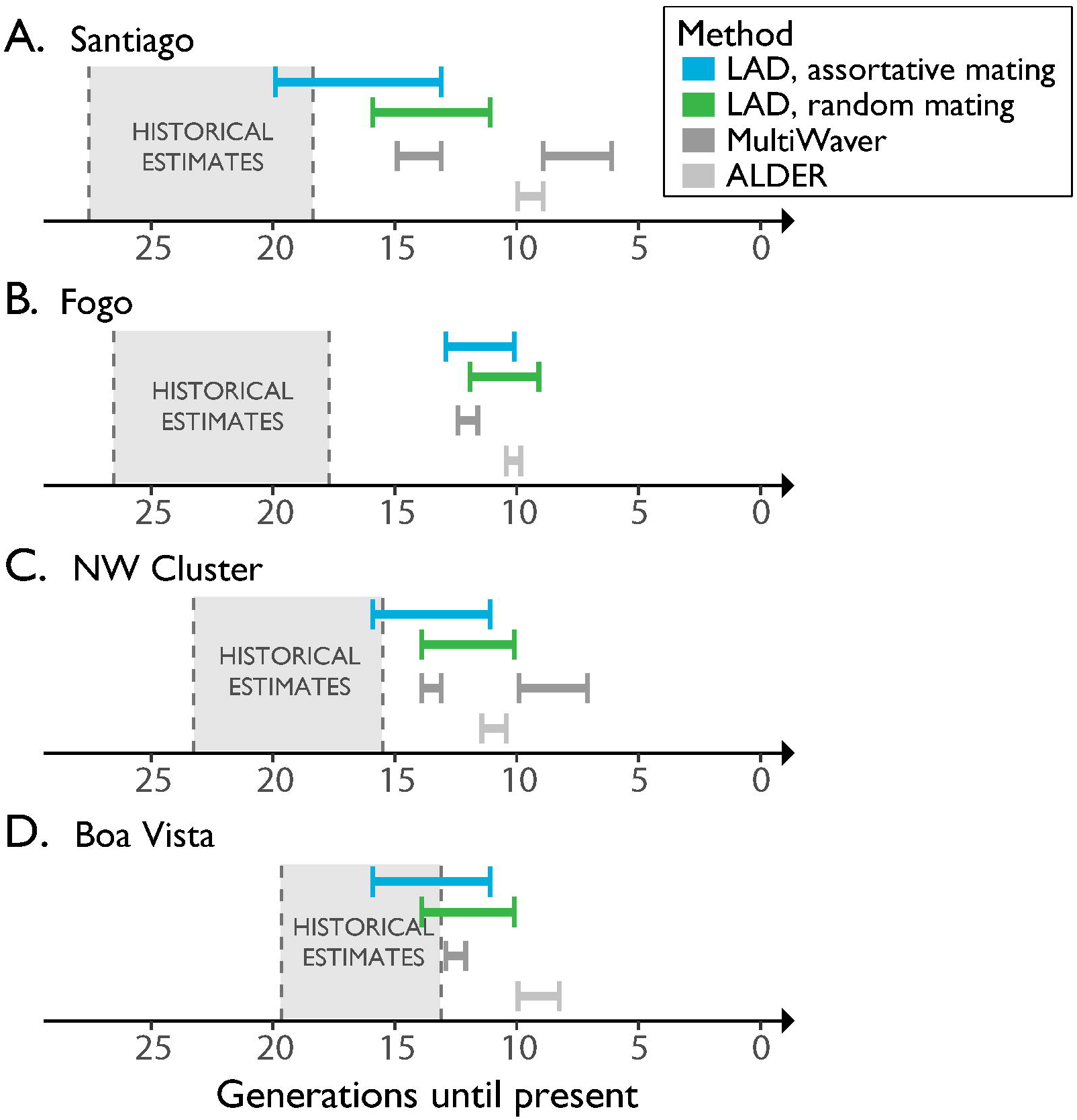
Timelines of inferred generations of admixture for each island of Cabo Verde. The island regions are listed in order of known settlement timing beginning with Santiago (A), followed by Fogo (B), and then the later migrations to the Northwest Cluster (C) and Boa Vista (D). For each island, admixture timing inferred with population genetic methods are shown in comparison to historical records. Historical estimates of when each island was first settled are shown in generations, based on a range of generation times (20-30 year generation time). For the LAD-based estimates of admixture timing, we tested a range of ancestry-assortative mating strengths. Specifically, we tested an assortative mating strength of 0 (results shown in green) vs values inferred using ANCESTOR in Fig 2 (results shown in blue). Within each of these LAD-based estimates of admixture timing, the presented interval reflects the range of estimates generated under the assumption of constant migration rate (m = 0.01; yielding older estimates) to the assumption of no migration (m = 0; yielding more recent estimates). MultiWaver results show the admixture generations inferred under the admixture model selected by MultiWaver, with 0.95 CI.

We additionally applied two strategies, ALDER (Loh et al. 2013) and MultiWaver 2.0 (Ni et al. 2019), that are not intended to account for assortative mating. ALDER uses the extent of LD decay among neighboring loci to infer mixture proportions and dates of a single admixture event (i.e., under an isolation model). In contrast, MultiWaver uses the length distribution of ancestral tracks to infer the best-fitting discrete (isolation or multi-wave) or continuous admixture model and then estimates the admixture parameters under the considered model. Assuming a generation time of 25 years, decay rates estimated with ALDER suggest admixture timing in the mid to late 1700s (Fig 3; all timing estimates are shown in Supp Table 1).

MultiWaver chose a multi-wave admixture model for Santiago (Fig 3A) and the Northwest Cluster (Fig 3C), but selected an isolation model for Fogo (Fig 3B) and Boa Vista (Fig 3D).

The estimates of admixture timing that do not explicitly consider ancestry-assortative mating, from ALDER and MultiWaver, are noticeably more recent than the known founding of Cabo Verde in late 1400s. Results from the LAD-based method described in Zaitlen et al. (2017) (Zaitlen et al. 2017) align more closely with historical records, particularly when we allowed for both ancestry-assortative mating and constant migration (Fig 3, Supp Fig 8). While multiple complex demographic factors likely impact these estimates of admixture timing (see “Patterns of genetic variation and admixture in Cabo Verde influence demographic inference” in the Discussion), the evidence of ancestry-assortative mating in Cabo Verde (Fig 2, Supp Fig 7) and the LAD-based inferences incorporating assortative mating (Fig 3, Supp Fig 8) suggest that explicitly accounting for ancestry-assortative mating improves estimates of admixture timing, while the assumption of random mating leads to underestimation of the age of admixture.

### Runs-of-homozygosity (ROH) reflect the contributions of the source populations and patterns of nonrandom mating

ROH arise when IBD haplotypes are inherited from a common ancestor. Thus, distinct length classes generally reflect inheritance from ancestors at different timescales in the population’s history. However, analyzing ROH of different length classes can be sensitive to arbitrary decision making in the classification process. To characterize long vs short ROH, we identified ROH using several approaches, beginning with the method of Pemberton et al. (2012) as implemented in the software GARLIC (Pemberton et al. 2012; Szpiech et al. 2017). This method classifies ROH into three categories (short, medium, and long) based on the modeling of the length distribution in each population as a mixture of Gaussian distributions. We also classified ROH based on the commonly used rule of thumb that a time depth of *m* meioses is expected to give a mean IBD segment length of 100/*m* cM (Thompson 2013). In this case, common ancestors within the past 20 generations (post-admixture in Cabo Verde) would give an expected IBD segment length of approximately 2.5 Mb or longer, so we applied 2.5 Mb as the shorter/long ROH boundary. This approximation uses the commonly applied baseline recombination rate of 1 cM/Mb, though we note that recombination varies among populations and along genomes (Hassan et al. 2021). Under these two classification schemes, we examined the distributions of total ROH per genome (sum of ROH segments of a specific length) within Cabo Verde, partitioning the distributions into length classes that are enriched for pre- and post-colonization processes. Under the ROH classification model of GARLIC, short and medium ROH formed due to processes that largely occurred within the source populations prior to the founding of Cabo Verde. Thus, while ROH in non-admixed populations are often modeled as three length classes, we emphasize the differences in pre- and post-admixture dynamics in Cabo Verde by focusing on shorter ROH (comprising both short and medium as called with GARLIC) vs. long ROH (Fig 4A). We observed similar ROH distributions using the shorter/long length class boundary of 2.5 Mb, based on the number of admixture generations (Fig 4B). To further support that ROH trends are concordant across classification schemes, we repeated this analysis under three additional length classification schemes, using empirically-based length class boundaries from the literature (see Methods; Supp Fig 9).

**Fig 4.**
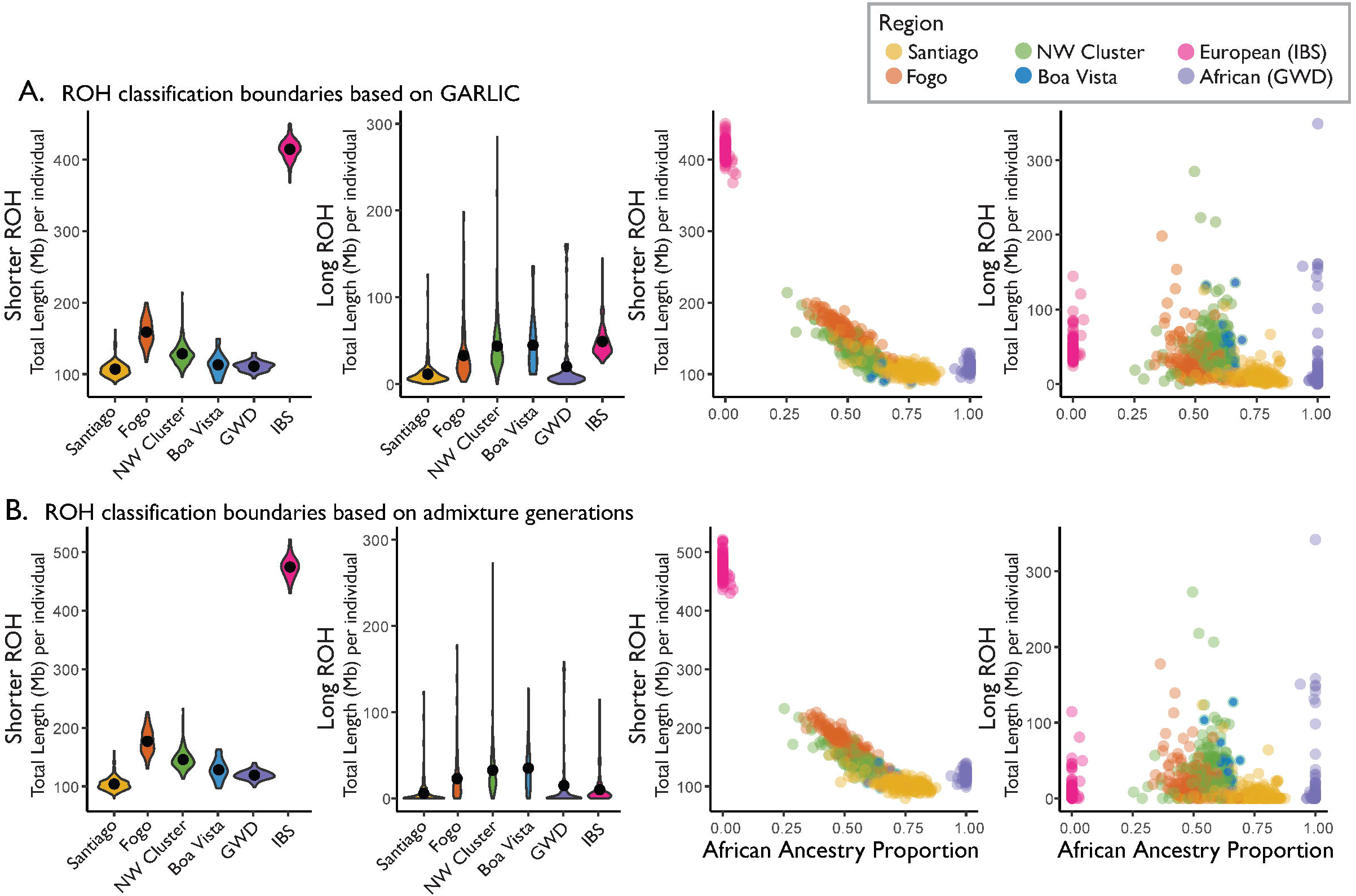
ROH content by island and in the context of ancestry. The total length of ROH per individual genome is plotted under (A) GARLIC-inferred ROH class boundaries (shorter/long ROH boundary of approximately 1.08 Mb for Santiago, 1.08 Mb for NW Cluster, 1.12 Mb for Fogo, 1.14 Mb for Boa Vista, 1.26 Mb for West African reference individuals, and 0.91 Mb for European reference individuals); and (B) under class boundaries estimated based on the number of admixture generations (shorter/long ROH boundary of 2.5 Mb). Violin plots show the population-specific distributions of the total (summed over each genome) length of autosomal ROH per individual. Solid black dots represent the within-population means. Scatter plots show the total length of autosomal ROH per individual plotted against West African ancestry proportions and colored by population.

ROH patterns in admixed populations should be interpreted with caution, since methods for detecting and interpreting ROH have been developed primarily in non-admixed populations (Pemberton et al. 2012; Ceballos et al. 2018). Thus, we consider ROH in Cabo Verde as an example of ROH distributions in admixed populations where we can explore results in the context of ancestry and population history. For example, to test for genetic evidence that population bottlenecks contributed to observed patterns, we estimated ancestry-specific population sizes (Supp Fig 10). Ancestry-specific population sizes in Cabo Verde suggest bottlenecks of both source populations, followed by population expansion within the past 10 generations (Browning and Browning 2015; Browning et al. 2018). If the length of ROH in an individual’s genome is driven by distance from Africa, as expected based on previous work (McQuillan et al. 2008; Pemberton et al. 2012), we predict the total length of ROH per genome to negatively correlate with West African admixture proportion. Consistent with this expectation, we observe a negative correlation between the total ROH and West African ancestry proportions in Santiago, Fogo, and the Northwest Cluster (Supp Fig 11A; correlations and significance values for total ROH and by length class are provided in Supp Fig 11). Breaking total ROH into shorter and long ROH, we see that this negative correlation is driven primarily by shorter ROH (Fig 4; Supp Fig 11B-C, Supp Fig 9).

Comparing the Cabo Verdean islands to each other, we observed the lowest levels of total ROH in all length classes of ROH in Santiago (pairwise Mann-Whitney U tests p < 1 × 10^−8^). The low amounts of all length classes of ROH in Santiago are consistent with several historical attributes of the islands, including serial founding beginning with the settlement of Santiago. Serial founding resulting in bottlenecks is expected to increase shorter ROH by reducing the number of ancestral lineages. But because shorter ROH reflects background relatedness from the source populations, shorter ROH are expected to be more sensitive to the contributions of the source populations.

Despite recent colonization bottlenecks, we observed that some admixed individuals in Cabo Verde present lower levels of ROH per genome than either of the source population proxies. Specifically, our classification of ROH into shorter and long ROH (Fig 4; Supp Fig 9) revealed that Santiago has significantly lower levels of shorter ROH compared the West African reference population (Mann-Whitney U p = 5.87 × 10^−7^), while long ROH are not depleted (Mann-Whitney U p = 0.841). This observation that Santiago has significantly lower levels of shorter ROH compared to even the West African reference is perhaps surprising given the canonical view of African source populations as generically having the lowest ROH in worldwide samples. This is consistent with the idea that shorter ROH tend to come from older shared ancestors in the source populations, and that these tracts can be further interrupted with local ancestry from another source population upon admixture. In contrast, long ROH are more likely to be driven by post-admixture dynamics such as small population size and mating preferences, and long ROH tracts may span multiple local ancestries. Indeed, we observed that ROH often span multiple ancestries, with long ROH spanning more switches in local ancestry per megabase. Specifically, 1.21% of shorter ROH contained at least one ancestry switch and 18.99% of long ROH contained at least one ancestry switch. Dividing ancestry switches by tract length, shorter ROH contain an average of 0.072 ancestry switches/Mb (standard deviation = 0.89), while long ROH contain an average of 0.089 switches/Mb (standard deviation = 0.24) (pairwise Mann-Whitney U test p < 1 × 10^−8^).

While ROH patterns in Cabo Verde are consistent with several historical observations, we again note that caution is warranted in applying methods and expectations built using non-admixed populations. We also note that, though we used a model-based ROH detection approach that included steps to mitigate genotyping errors and biases in allele frequency estimates, sensitivity and specificity of ROH detection is generally lower for shorter ROH. Further, records suggest that enslaved African individuals in Cabo Verde came from the Senegambian region of Africa, and lack of genomic data from this vast region makes it difficult to assess how the genetic diversity of contributing African peoples might contribute to low shorter ROH levels in Cabo Verde. These sources of uncertainty necessitate caution in inferring population history from ROH distributions. However, the observed differences in ROH among the islands suggest that ROH are sensitive to population genetic processes even on the short timescale since Cabo Verde’s settlement. Together, these observations and caveats underscore the need for future work testing the effects of admixture on ROH and on ROH detection, as we explore in the Discussion.

### Ancestry patterns on the X chromosome vs the autosomes reflect sex-biased demographic processes

To further understand population structure in Cabo Verde, we next consider how autosomal and X-chromosome ancestry and genetic variation patterns can be used to infer the sex-specific demographic history of the last ∼20 generations since founding. Some earlier analyses of post-admixture population structure using Y-chromosome diversity suggested sex-biased demographic processes (Beleza et al. 2012). Here, we examine autosomal and X-chromosome ancestry to test for sex-biased migration. On all islands, autosomal versus X chromosome ancestry patterns suggest that male and female contributions differ significantly by source population (Fig 5A). Specifically, there is higher West African ancestry on the X chromosome than the autosomes (mean African ancestry proportion for the X chromosome = 0.76; mean African ancestry proportion for autosomes = 0.60; Wilcoxon Signed-Rank Test p < 1 × 10^−8^ for Santiago, Fogo, and the Northwest Cluster; p = 0.0013 for Boa Vista), consistent with sex-biased contributions of the founders. To quantitatively explore these differences in male vs female demographic history, we used a model-based approach to estimate differences between male and female contributions (Goldberg and Rosenberg 2015). A model of constant admixture is supported over a model of instantaneous admixture by historical records (see Methods for historical data sources) and by our comparison of admixture timing estimates (above). Using the method of Goldberg and Rosenberg for a model of constant admixture, we compared observed and expected X-chromosomal and autosomal ancestry over a grid of possible parameter values. Following the implementation from Goldberg et al. (2017) (Goldberg et al. 2017), Fig 5B presents results from the 0.1% of parameter sets closest to observed data based on a Euclidean distance between observed population mean ancestries on the X and autosomes and expected ancestries under the model (Supp Fig 12 shows the same data represented by plotting the male contributions). These estimated sex-specific contributions from West African and European source populations under a model of constant admixture suggest greater contributions from West African females and European males.

**Fig 5.**
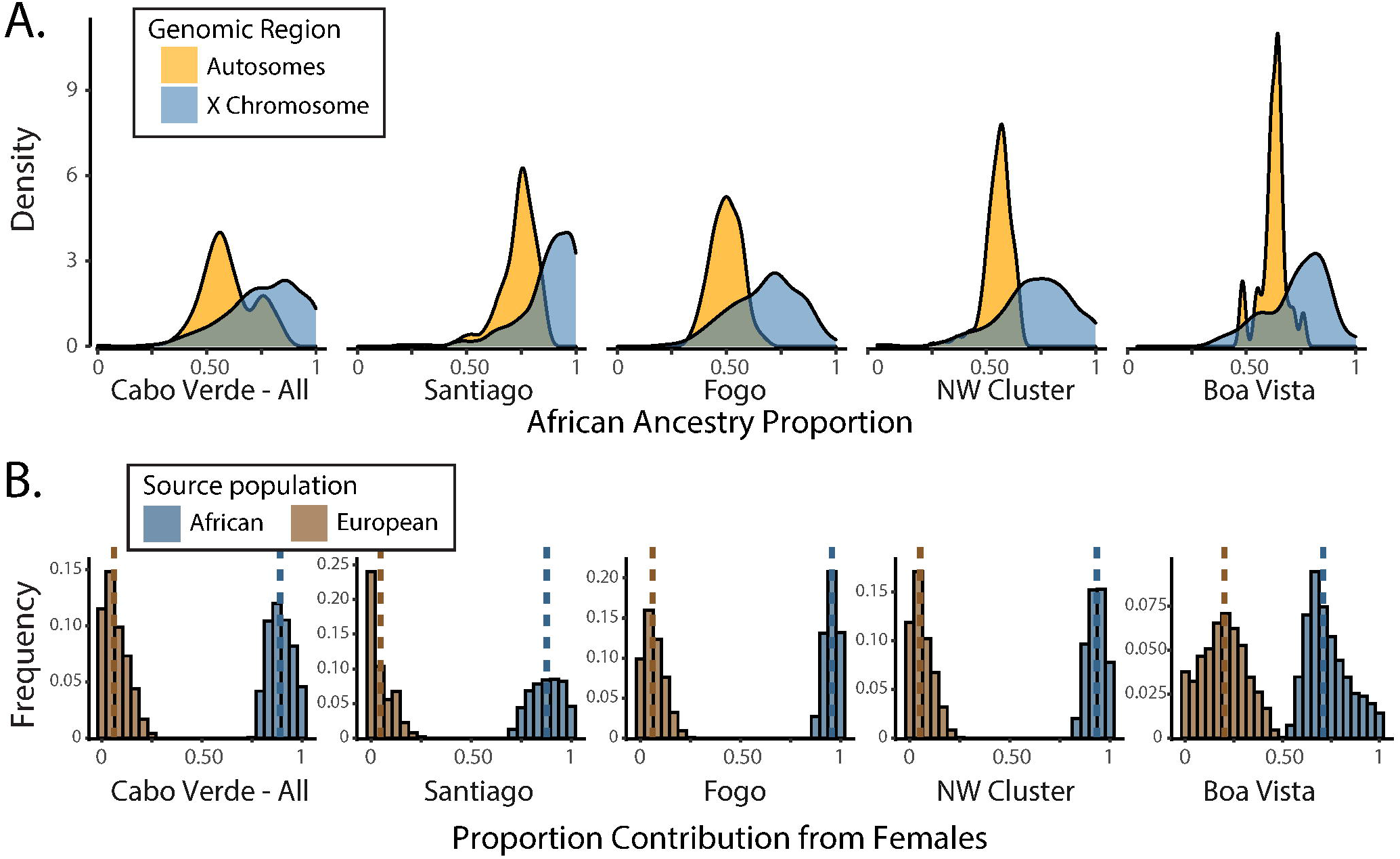
Sex-biased admixture in Cabo Verde. (A) The distribution of West African ancestry proportion on the autosomes and X chromosome for each of the island regions, estimated with ADMIXTURE. (B) Under a model of constant admixture over time, the fraction of the total contribution of genetic material originating from females for West African and European source populations. Here, we show the distribution of parameter sets for the smallest 0.1% of Euclidean distances between the model-predicted and observed X and autosomal ancestry from a grid of possible parameter values. The range of sex-specific contributions from West African and European source populations that produce ancestry estimates closest to those observed in Cabo Verde are shown for Cabo Verde as a whole (left), and then broken down by region, with medians (dashed lines).

Given our observation that source populations make distinct contributions to ROH in Cabo Verde (Fig 4), we next investigate differences in autosomal vs X chromosome ROH content. Differences in autosomal vs X chromosome ROH may arise in part due to the source populations contributing in a sex-biased manner. Notably, this effect would be seen most in shorter ROH, since shorter ROH reflect the homozygosity of older haplotypes and background relatedness from the source populations. However, it is challenging to disentangle processes shaping X vs autosomal ROH, due to the smaller effective population size of the X chromosome. Our results suggest that sex-biased admixture processes in Cabo Verde are reflected in ROH, with lower levels of shorter ROH on the X chromosome than on autosomes (Fig 6A). European individuals have higher levels of ROH than West African individuals, and the higher contributions of African X chromosomes (vs European X chromosomes) may drive the lower levels of short ROH in Cabo Verde on the X chromosome vs the autosomes (Fig 6A). Long ROH reflects different dynamics, potentially including the reduced post-admixture population size and other sex-specific processes.

**Fig 6.**
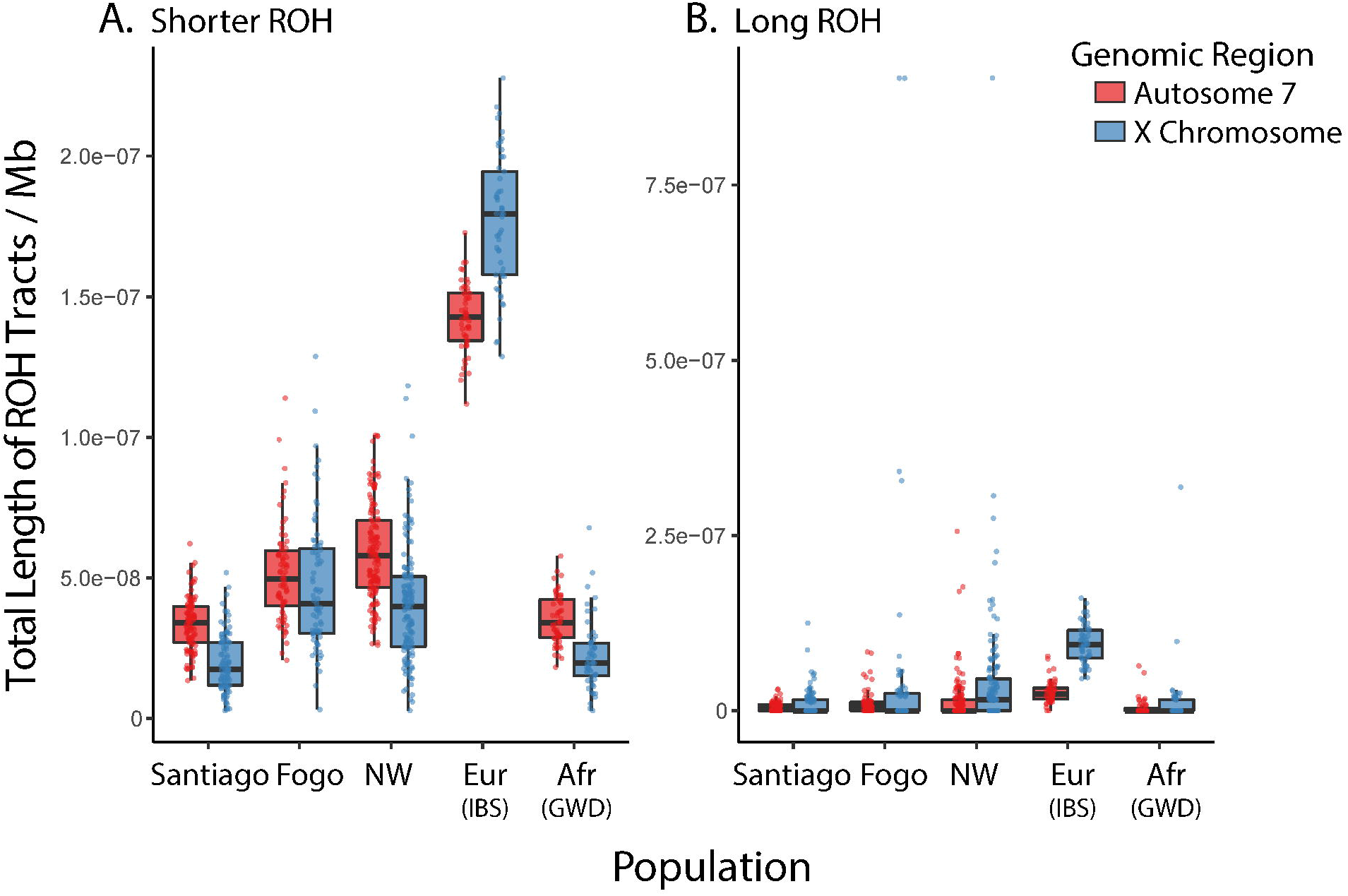
Autosomal (chromosome 7) vs X chromosome distributions of ROH by class. (A) shows the population-specific distributions of the total length of shorter ROH per individual on the X chromosome compared to an autosome. Chromosome 7 was chosen as the autosomal point of comparison, given that it is the autosome most similar in size to the X chromosome. Totals are shown for females only, so that the same samples are being compared across the X chromosome and chromosome 7. Totals are plotted per Mb to account for slight differences in chromosome lengths. (B) shows the same information for long ROH.

## Discussion

In this study, we leveraged patterns of genetic variation in Cabo Verde to infer the demographic history of the past ∼20 generations. We found that distinct genetic patterns of four island regions within the archipelago reflect the colonization history of the islands, including island-specific settlement timing, admixture dynamics, mating patterns, and sex-biased demography. Together these results demonstrate how patterns of ancestry and genetic variation are shaped by social and demographic forces on short timescales. By better understanding how complex population histories generate genetic variation, we can improve interpretation of inference from populations without historical records.

### Patterns of genetic variation and admixture in Cabo Verde reflect colonization history and subsequent sociocultural dynamics

The observed island-specific genetic patterns reflect the complex ecological and social factors that shaped the settlement of Cabo Verde. Historical evidence suggests that the islands were founded in a stepping-stone pattern, with settlement stages that were accompanied by important economic and sociocultural shifts (Correia e Silva 2002). Early written records of the population (described below) suggest that admixture began early during the settlement of Cabo Verde, despite the highly racially stratified, slavery-based system that characterized the first settlement stage (Cabral 2001). The second and third stages were carried out by mostly already-admixed individuals who had become a significant group within Cabo Verdean society and who migrated from the southern to the northern islands (Correia e Silva 2001). We found that the staggered settlement history and the island-specific population dynamics shaped patterns of ancestry and genetic variation within Cabo Verde.

Admixture in Cabo Verde is consistent with a model of continuous gene flow from Europe and West Africa, with European males and African females contributing predominantly to the Cabo Verdean gene pool, as seen in other admixed populations that result from the trans-Atlantic slave trade (Eltis et al. 2010; Micheletti et al. 2020). PCA (Supp Fig 1), IBD (Fig 1, Supp Fig 3-5), and kinship (Supp Fig 6) are consistent with genetic drift occurring during the archipelago’s settlement history of consecutive founder effects and subsequent relative isolation of the islands. Specifically, individuals are genetically more related within an island than among islands, and considerably more individuals from the northern islands and Fogo are connected to each other by high IBD pairwise connections (Supp Fig 4-5). We observed differences in IBD that reflect the contributions of the source populations, such as Santiago having both the highest West African ancestry proportions and the lowest overall levels of IBD sharing. The higher African genomic ancestry on Santiago is consistent with previous results (Beleza et al. 2012; Beleza et al. 2013), and indicates differences in settlement patterns despite the fact that colonization of the other islands involved migrations from Santiago and was based on the same slavery-based system.

Using a mechanistic model of sex-biased admixture (Goldberg and Rosenberg 2015), we used differences in ancestry on the X chromosome vs the autosomes to examine sex-biased migration in Cabo Verde. We found that admixture occurred primarily through the mating of European males and African females, consistent with historical work that documents sex bias during the settlement of Cabo Verde. First records of the demographic distribution in Cabo Verde are found in a letter by a judicial official Pero de Guimarães to the Portuguese King in 1513, which stated the presence of 118 European individuals in Santiago, of which only four were (single) women (Pero de Guimarães 1991). The marginal presence of European females throughout the founding of Cabo Verde contributed to the extensive genetic admixture currently found in the archipelago. The distribution of ROH in autosomes versus X chromosome is influenced by initial male versus female contributions to the admixed population’s gene pool. Throughout the islands, the higher contributions of African X chromosomes (vs European X chromosomes) may drive the lower levels of short ROH in Cabo Verde on the X chromosome vs the autosomes (Fig 6A). These results underscore the long-lasting genetic impacts of the trans-Atlantic slave trade, as has been recently shown in admixed populations across the Americas (Micheletti et al. 2020).

Despite only a few hundred years of unique population histories among the islands, we also found that patterns of genetic variation reflect island-specific patterns in post-admixture dynamics. Observed patterns of IBD, kinship, and long ROH are all consistent with overall high levels of relatedness within Cabo Verde, particularly in Fogo and the northwestern islands. In other worldwide populations, similarly high IBD levels have been attributed to founder effects (Campbell et al. 2012 Jul 31; Moreno-Estrada et al. 2013). In Cabo Verde, patterns of relatedness may be shaped by both founder effects and by post-admixture dynamics such as nonrandom mating keeping similar haplotypes together rather than distributing them randomly throughout mating pairs in the population. Indeed, we inferred a positive correlation between mating pairs of the previous generation, suggesting ancestry-assortative mating on the short timescale since the founding of the islands. This correlation between mating pairs was significant within all islands except for Boa Vista, though the inferred Pearson’s R in Boa Vista still differed significantly from the distribution of random samples of parental haplotypes, shown in Supp Fig 7. Thus, we expect that the lack of significance of Boa Vista’s correlation in parental ancestries in Fig 2 may simply be because the sample size was lowest for Boa Vista (n = 26). Nonetheless, we highlight differences in mating patterns among the islands as a topic that could be explored further in future studies. For example, our inferences suggest that assortative mating appears the strongest in Santiago. Santiago has many unique demographic features, such as being the oldest population and having the largest population size of the islands. As we learn more about ancestry-assortative mating, future investigations may shed light on how these demographic attributes and others contribute to mating patterns.

Mating patterns and sex-biased admixture are distinct—but interacting—processes in the history of Cabo Verde. Sex-biased migration determined the gene pool within Cabo Verde, while patterns of nonrandom mating within the islands shaped subsequent generations of admixture. These mating patterns were tied to social and economic structure. European enslavers sexually coerced enslaved African women during slavery. After the end of slavery, racially unbalanced control of landownership and higher social status of Europeans resulted in continued socioeconomic pressures for African women to have children with European men. Such unions often resulted in improved socioeconomic status for both the women and their children. For example, the first admixed individual to assume an important position within the administration of Santiago was recorded in 1546 (Seibert 2014 Jun 1). Over generations, Cabo Verde became a population of extensively admixed ancestry, with mating patterns shaped by the interconnected effects of social position and skin color. This type of nonrandom mating may have been more prominent in the second half of the 17^th^ century, due to the large exodus of the white population following the decay of the slave trade. With this population shift, historical sources say that the socioeconomic elite of the islands became predominately the “white children of the islands” (Cabral 2001), leading to further racial segregation. This historical evidence of nonrandom mating patterns—together with our genetic observations— demonstrates how quickly changes in social structure can impact population homozygosity and differentiation.

The island of Fogo stands out as having unique social and historical processes, even though it was settled shortly after Santiago. Fogo’s society was, since its origins, a conservative rural society whose main economic activity was to produce goods to trade in the African coast. The socioeconomic elite was patriarchal and aristocratic (Cabral 2012), and composed primarily of related families, which promoted first-cousin marriages (Barbosa, Gilda 1997). The sustenance of this class depended on land ownership, which allowed the slavery-based system to be perpetuated longer in Fogo than in Santiago (Correia e Silva 2002; Cabral 2012). The unique attributes of Fogo are reflected in population genetic patterns. For example, we found that African ancestry proportion align with kinship patterns overall, but exceptions were most obvious in individuals from Fogo and the Northwest Cluster, where some observed relatedness patterns are not as clearly explained by global ancestry proportions. In our estimation of admixture timing using local ancestry disequilibrium, we observed that Fogo has greater levels of local ancestry disequilibrium than the other islands. Higher local ancestry disequilibrium may be driven by ancestry-assortative mating. Additionally, it may be that Fogo has experienced stronger founder effects, which would decrease the number of ancestral lineages. Indeed, the higher levels of shorter ROH within Fogo (Fig 4; Supp Fig 9) are consistent with founder effects increasing background relatedness and thus increasing shorter ROH. Though we emphasize the need for further work to understand the dynamics of ROH in admixed populations, these patterns are in agreement with the lower Y-chromosome haplotype diversity observed in Fogo compared to the other islands (Beleza et al. 2012). Long ROH and IBD patterns gives us further insights about the post-admixture demographic processes in the archipelago, including uncovering genetic consequences of consecutive founder effects.

### Patterns of genetic variation and admixture in Cabo Verde influence demographic inference

The admixture dynamics, founder effects, and mating patterns within Cabo Verde shape summaries of genetic variation that are often used to inform demographic inference, such as local ancestry disequilibrium, IBD, and ROH. To investigate the effects of these population dynamics on demographic inference, we estimated admixture timing using multiple population genetic tools. Estimates of admixture timing from genetic data were most concordant with historical records when using inference based on local ancestry disequilibrium (Zaitlen et al. 2017), allowing for ancestry-assortative mating and migration after the founding of the admixed population. Under this method, we tested a range of ancestry-assortative mating strengths (0 vs values inferred using ANCESTOR in Fig 2 and a broader range of values in Supp Fig 8) and migration rates (0 vs 1% each generation). The migration levels are used to demonstrate the trends of timing estimates using constant migration and assortative mating, rather than to obtain a specific estimate of the migration parameters. Consistent with previous theoretical work on the impact of assortative mating on the timing of admixture, we find that estimates that do not account for ancestry assortment are more recent than historical records and than estimates that include ancestry assortment (Zaitlen et al. 2017; Goldberg et al. 2020). Under random mating, haplotypes are distributed randomly in mating pairs in a population, allowing recombination to shuffle haplotypes over generations. In contrast, ancestry-assortative mating can keep ancestral haplotypes more intact, leading to underestimation of the number of generations of admixture when estimating admixture timing. Zaitlen et al. (2017) and Goldberg et al. (2020) theoretically demonstrate that certain models of non-random mating in admixed populations drive variation in the ancestry proportion and maintain linkage over time— summary statistics that will make admixture appear more recent than it is when mating patterns are not considered (Zaitlen et al. 2017; Goldberg et al. 2020). Our evidence of ancestry-assortative mating in Cabo Verde underscores the importance of accounting for nonrandom mating in understanding admixture and inferring demographic history. Assortative mating has been documented with respect to genetic ancestry, socioeconomic factors, and phenotypic characteristics (Risch et al. 2009; James Y Zou et al. 2015; Robinson et al. 2017; Sebro et al. 2017; Zaitlen et al. 2017), but remains an under-recognized force in human population genetics and demographic inference.

Notably, admixture timing estimates were most concordant with the historical records when allowing both assortative mating and constant admixture. An admixture scenario that allows multi-wave or constant admixture model is compatible with historical work suggesting constant migration from both Europe and West African throughout the settlement of the islands. Importantly, the authors of ALDER and MultiWaver point out that methods for inferring admixture timing have varying sensitivities to different admixture scenarios (Loh et al. 2013; Ni et al. 2019). ALDER notes that multi-wave or continuous admixture can lead to more recent estimates of admixture timing, as these processes can lead to an increase of LD due to admixture in every generation when compared to the isolation model. Despite this caveat, ALDER is frequently applied in cases where admixture is not strictly instantaneous.

MultiWaver inferred more complex admixture models for two out of the four island regions, but the consideration of additional migration parameters in the calculation of the admixture times led to results similar to our LAD model with random mating. We also note that MultiWaver may not be able to accurately infer demographic histories that deviate from its pre-defined admixture models. In general, all available demographic inference methods must make simplifying assumptions about demographic history. Our findings suggest that two simplifying assumptions (random mating and single-pulse admixture) push inferred estimates of admixture timing to appear more recent than the historical estimates.

While allowing both assortative mating and constant admixture yielded admixture timing estimates that were most consistent with the historical records, it is important to note that none of the inferred ranges of admixture timing captured the historically-documented onset of admixture. This may be due to additional simplifying assumptions underlying the demographic inference methods, such assumptions about the stationary of effective population size, the composition of the sample (i.e., that the sampled individuals are a random sample from the population), or the neutrality of evolution. All of the timing methods we used placed admixture timing for the different islands closer together than historical dates of settlement, consistent with historical expectations that the initial admixture in the southern islands was significant, and that many individuals that occupied the northern and eastern islands during the second and third settlement stages of Cabo Verde were already admixed. For example, the serial founding of the islands may explain why estimates of admixture timing for Boa Vista were closer to historical records. While Boa Vista was founded most recently, it was founded by already admixed individuals. This observation, together with IBD and kinship patterns, support the serial founding of the groups of islands as the main model of settlement of Cabo Verde, as opposed to their independent settling as some historical data suggest (Correia e Silva 2002). This type of serial founding scenario is common throughout recent human migration, underscoring that settlement patterns, in addition to settlement timing, are critical components of accurately inferring human demographic history.

We found that Cabo Verde’s island-specific demographic history and admixture dynamics have important genomic consequences, as observed with ROH. Despite the relatively recent colonization of the islands, some individuals presented even lower overall levels of ROH than African reference populations. We found that low overall levels of ROH in Cabo Verde are driven by shorter length ROH. This observation is consistent with the idea that shorter ROH can be attributed to older shared ancestors from the source populations, and these tracts can be interrupted with admixed ancestry upon admixture. However, many admixed individuals still present excess long ROH, likely reflecting post-admixture processes such as serial founding and ancestry-assortative mating. The observed island-specific distributions of ROH are consistent with the colonization history of the islands. For example, the cultural dynamics and higher rates of first-cousin marriages in Fogo (Barbosa, Gilda 1997), possibly driving high levels of long ROH (Fig 4). In contrast, Santiago has both the oldest population and the largest population size, and has comparatively low levels of both shorter and long ROH. These observations suggest that more work, both empirical and theoretical, is needed to understand the interacting forces of local ancestry and ROH.

In sum, we provide insights into the population history of Cabo Verde and demonstrate how admixed populations can provide powerful test cases for understanding demographic processes and genomic consequences in recent human history. We show that patterns of shared ancestry between and within the islands (quantified with IBD and kinship estimates) reflect serial founder effects as well as settlement patterns such as post-admixture nonrandom mating. We find that accounting for nonrandom mating allows us to improve inference of admixture timing and better contextualize genomic consequences of admixture dynamics, such as ROH. We find that differences in ancestry on the X chromosome vs the autosomes reflect sex-biased demographic processes. Given the ubiquity of admixture throughout modern human population, these results provide important, generalizable considerations for the study of recent human evolution.

## Data and Code Availability

The de-identified local ancestry calls, ROH calls, and IBD calls will be made publicly available by the time of publication. Inferred local ancestry information can be found at https://doi.org/10.5281/zenodo.4021277, and inferred ROH and IBD tracts can be found at [link/DOI to be inserted by the time of publication]. Code generated for this study can be found at https://github.com/agoldberglab/CaboVerde_Demographic_Analyses. Sampling consent forms do not allow for public release of genotype data, which were provided by Beleza et al. (2013). However, the original genotype data will be made available upon signing a material transfer agreement assuring that the data will only be used in accordance with the restrictions of the informed consent, and agreeing that the data will be destroyed after the research is complete.

## Acknowledgements

We thank Jorge Rocha for his contribution during project conception and sampling, and Greg Barsh for his support in later stages of data collection. We thank Noah Zaitlen for sharing scripts to assist in our application of the Zaitlen et al. (2017) local ancestry disequilibrium method. This work was supported by MRC-UK MR/M01987X/1 and FCT-Portugal PTDC/BIA-BDE/64044/2006 awarded to Sandra Beleza; NIH NIGMS grant R35 GM133481 awarded to Amy Goldberg; NIH NIGMS grant F32 GM139313 awarded to Katharine Korunes; and by CNPq (CSF/PDE 207651/2014-0) awarded to Giordano Bruno Soares-Souza. We acknowledge the source of the reference datasets: 1000G 30x data generated at the New York Genome Center with funds provided by NHGRI Grant 3UM1HG008901-03S1. We thank Drs. Ryan Hernandez, Jeffrey Ross-Ibarra, Kirk Lohmueller, Brenna Henn, and the anonymous reviewers for their helpful feedback on this work. Finally, we would like to thank the Cabo Verdean participants for their invaluable contributions, and the University of Cabo Verde administration for their support.

