## Supplementary Figures and Tables for "Sex-biased admixture and assortative mating shape genetic variation and influence demographic inference in admixed Cabo Verdeans"

**Supplementary Figures**

Supp Fig 1: Principal Components Analysis and estimated admixture proportions.
Supp Fig 2: Comparison to previous global and local ancestry inference.
Supp Fig 3: Total IBD between and within the islands in the context of geography.
Supp Fig 4: IBD by island.
Supp Fig 5: The distribution of IBD sharing within and between islands.
Supp Fig 6: Kinship estimation in the context of ancestry.
Supp Fig 7: Comparison of inferred ancestry-assortative mating strength to the sampling distribution based on randomly paired haplotypes.
Supp Fig 8: Timelines of inferred generations of admixture using LAD under varying degrees of ancestry-assortative mating.
Supp Fig 9: ROH distributions under various length classification cutoffs.
Supp Fig 10: Ancestry-specific population size.
Supp Fig 11: ROH vs African ancestry proportion.
Supp Fig 12: Sex-biased admixture in Cabo Verde – male contributions.

**Supplementary Tables**

Supp Table 1: Summary of computational methods.
Supp Table 2: ROH length classification and LOD score cutoffs from Garlic.
Supp Table 3: Estimates of admixture timing.

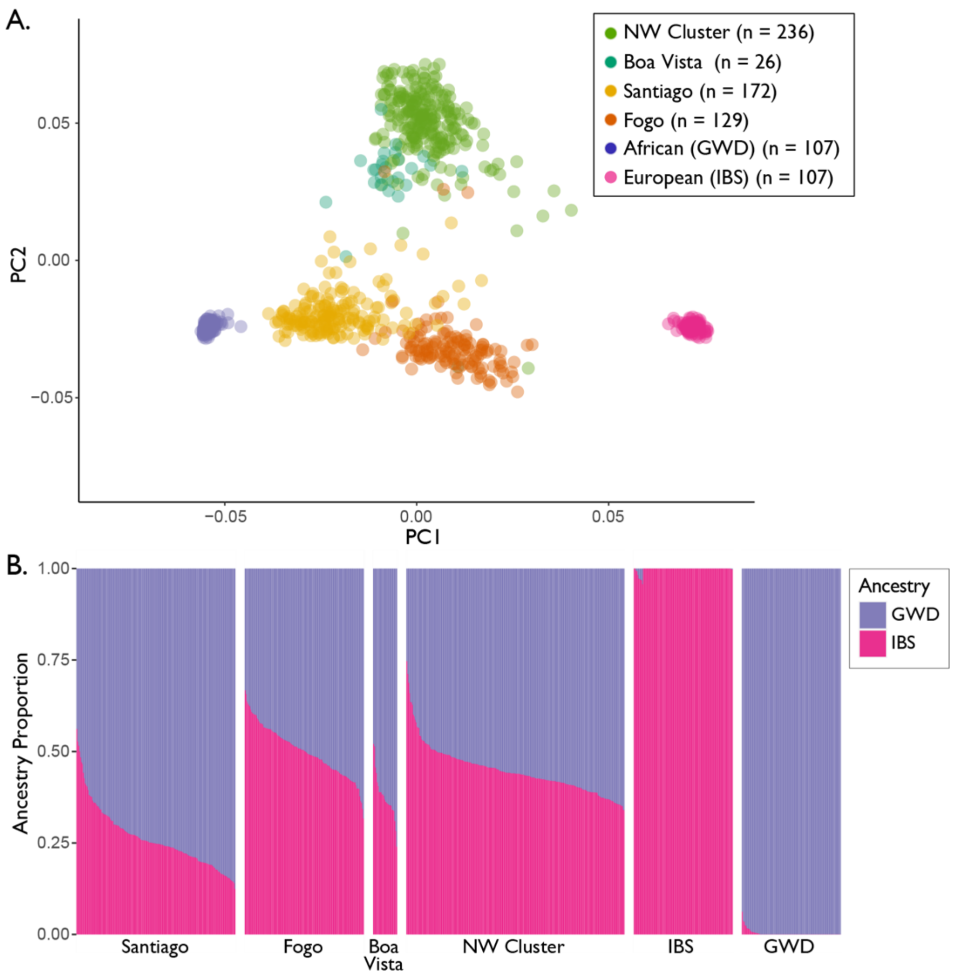


**Supp Fig 1: Principal Components Analysis and estimated admixture proportions**. (A) The first two PCs are shown, based on the pruned autosomal dataset of 514,551 SNPs from Cabo Verdean individuals (colored based their island of origin) and reference populations (West African and European), with sample sizes noted in the legend. (B) ADMIXTURE estimates of overall autosomal ancestry per individual, grouped by island region or reference population.

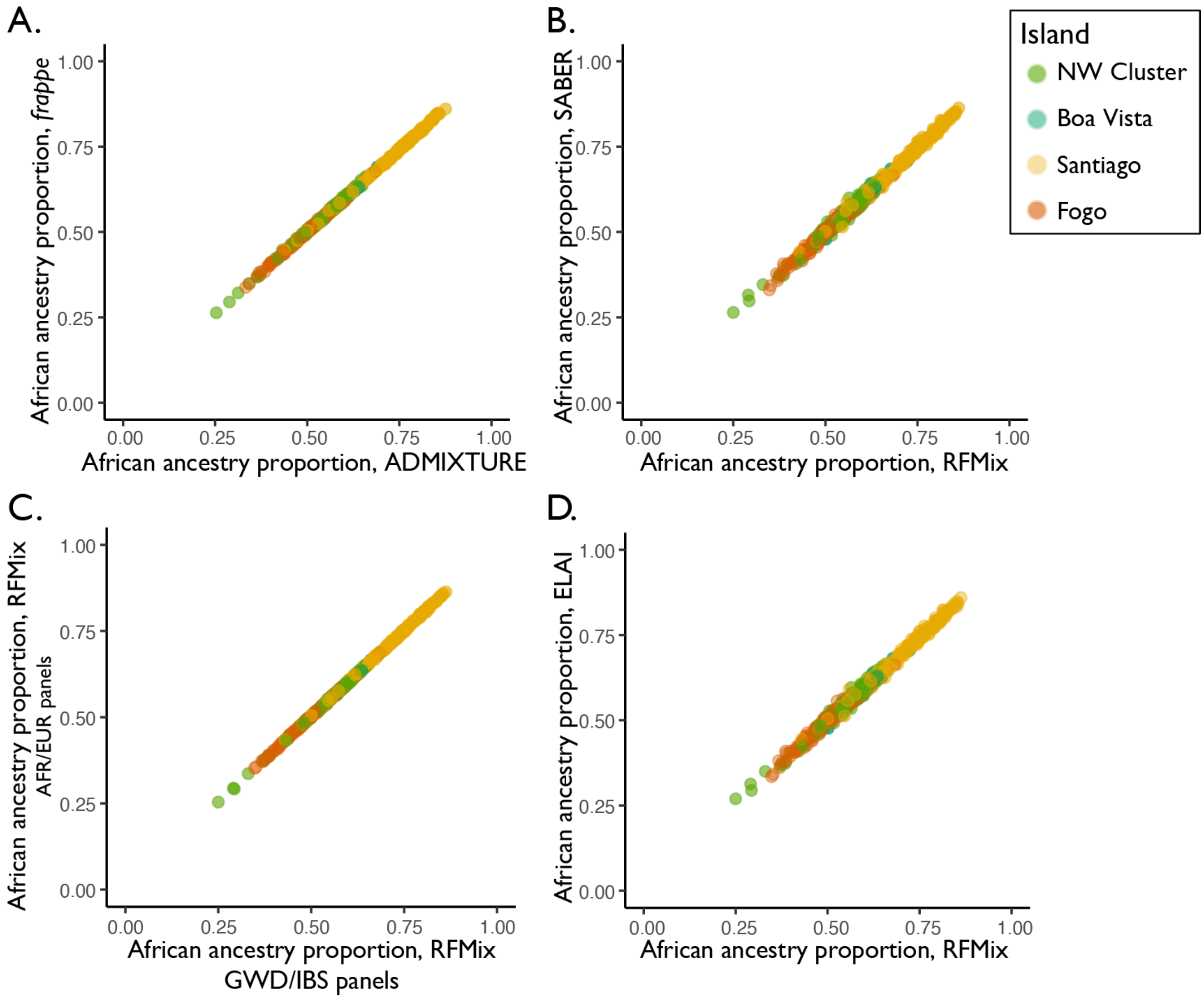


**Supp Fig 2: Comparison to previous global and local ancestry inference**. (A) ADMIXTURE estimates of overall autosomal ancestry per individual from this study are highly correlated (Pearson's $R > 0.99$, $p < 1 \times 10^{-8}$) with *frappe* estimates from Beleza et al. (2013). (B) Averaged autosomal ancestry per individual from local ancestry calls using RFMix are highly correlated (Pearson's $R > 0.99$, $p < 1 \times 10^{-8}$) with estimates from SABER local ancestry calls from Beleza et al. (2013). (C) RFMix local ancestry calling using 1kG resequenced genomes from all AFR and EUR populations correlates closely with RFMix local ancestry calling using GWD and IBS reference panels (Pearson's $R > 0.99$, $p < 1 \times 10^{-8}$). (D) Averaged autosomal ancestry per individual from local ancestry calls using RFMix correlate closely (Pearson's $R > 0.99$, $p < 1 \times 10^{-8}$) with those from ELAI.

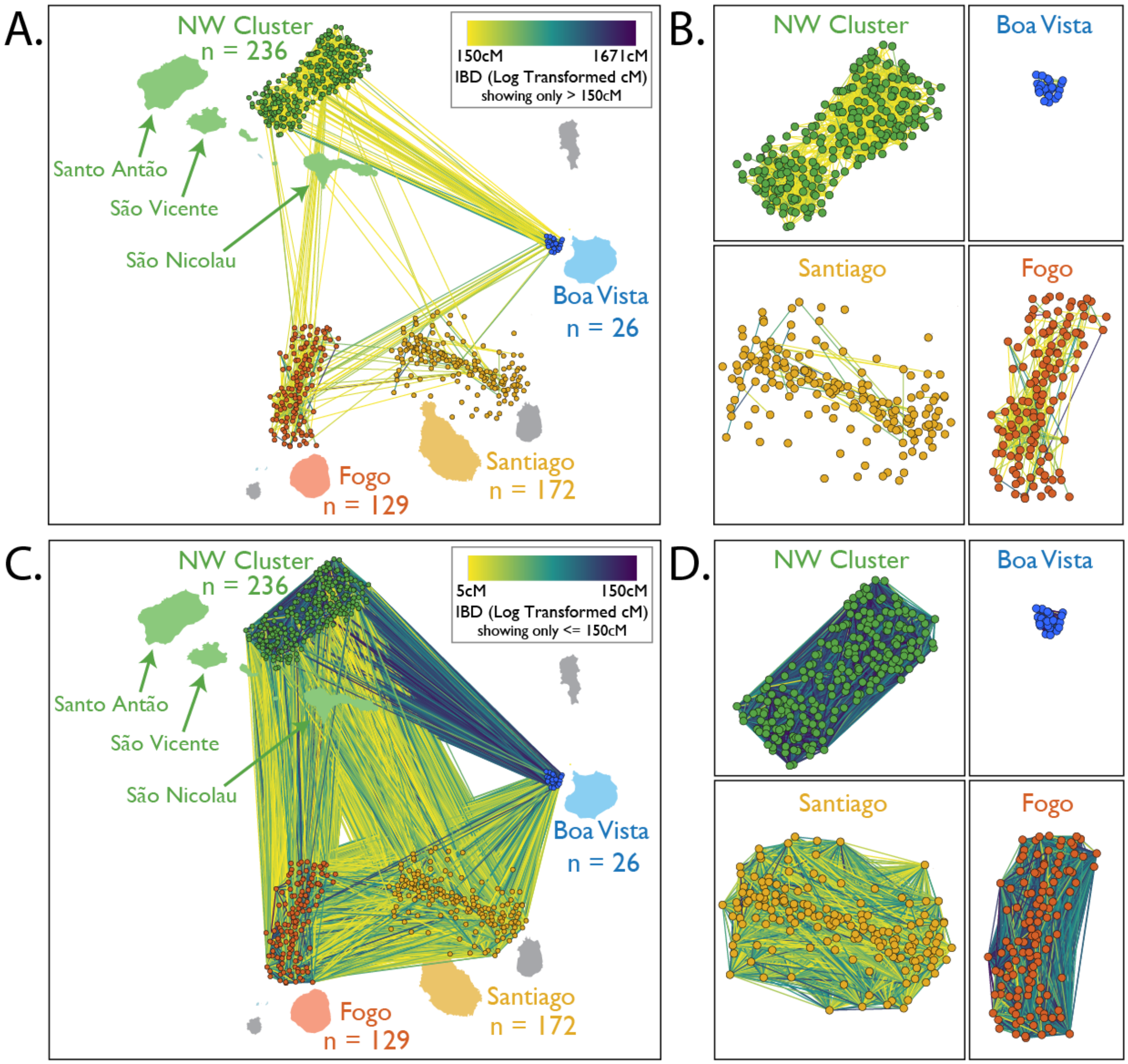


**Supp Fig 3: Total IBD between and within the islands in the context of geography.** The total length of segments identical-by-descent (IBD) are summed for each pairwise comparison of individuals. Here, we separately plot total IBD lengths over 150 cM (A and B) and less than or equal to 150 cM (C and D). In (A) and (C), each island has a corresponding cluster of nodes representing all sampled individuals. The edges between the nodes represent total IBD tract length shared between a pair of individuals. Individuals are localized to be adjacent to the island where they were sampled, and edges within an island's cluster attract nodes proportionally to shared IBD. Node placement within islands by a force-directed algorithm means that the spread of each cluster reflects the level of relatedness in each population. (B) and (D) show only the within-island IBD of (A) and (C), respectively. Note that (A) is the same as Figure 1, shown here for ease of comparison to within-island IBD.

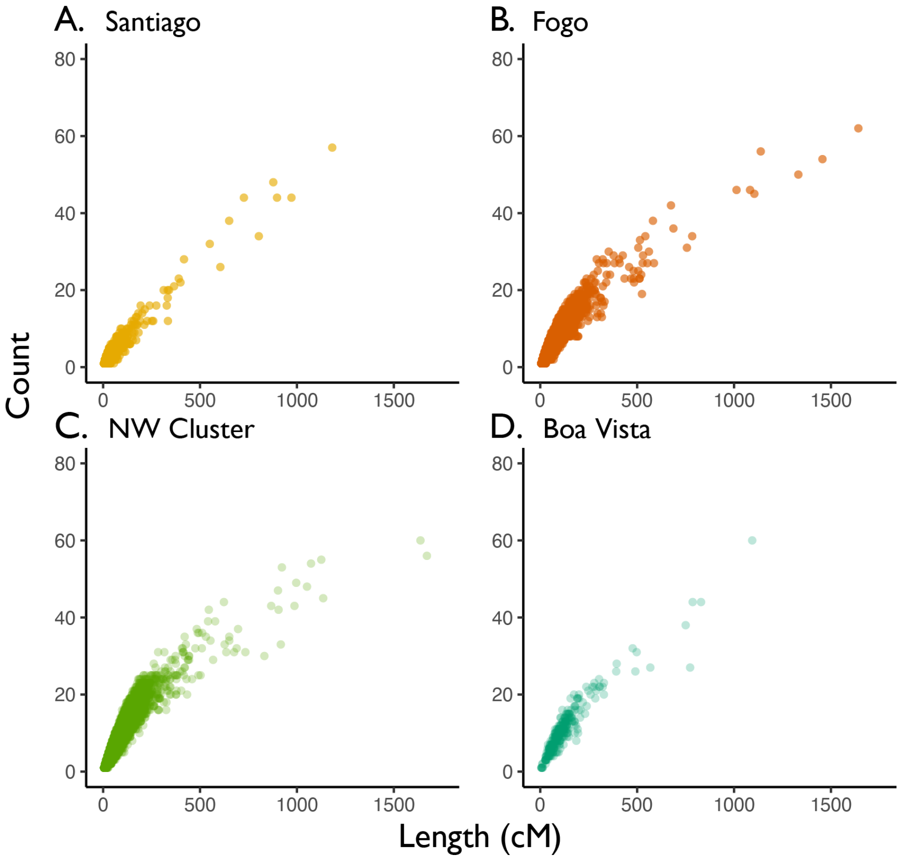


**Supp Fig 4: IBD by island.** Pairwise IBD sharing within islands. The total count and summed length of pairwise inferred IBD segments between individuals within each of four Cabo Verdean islands.

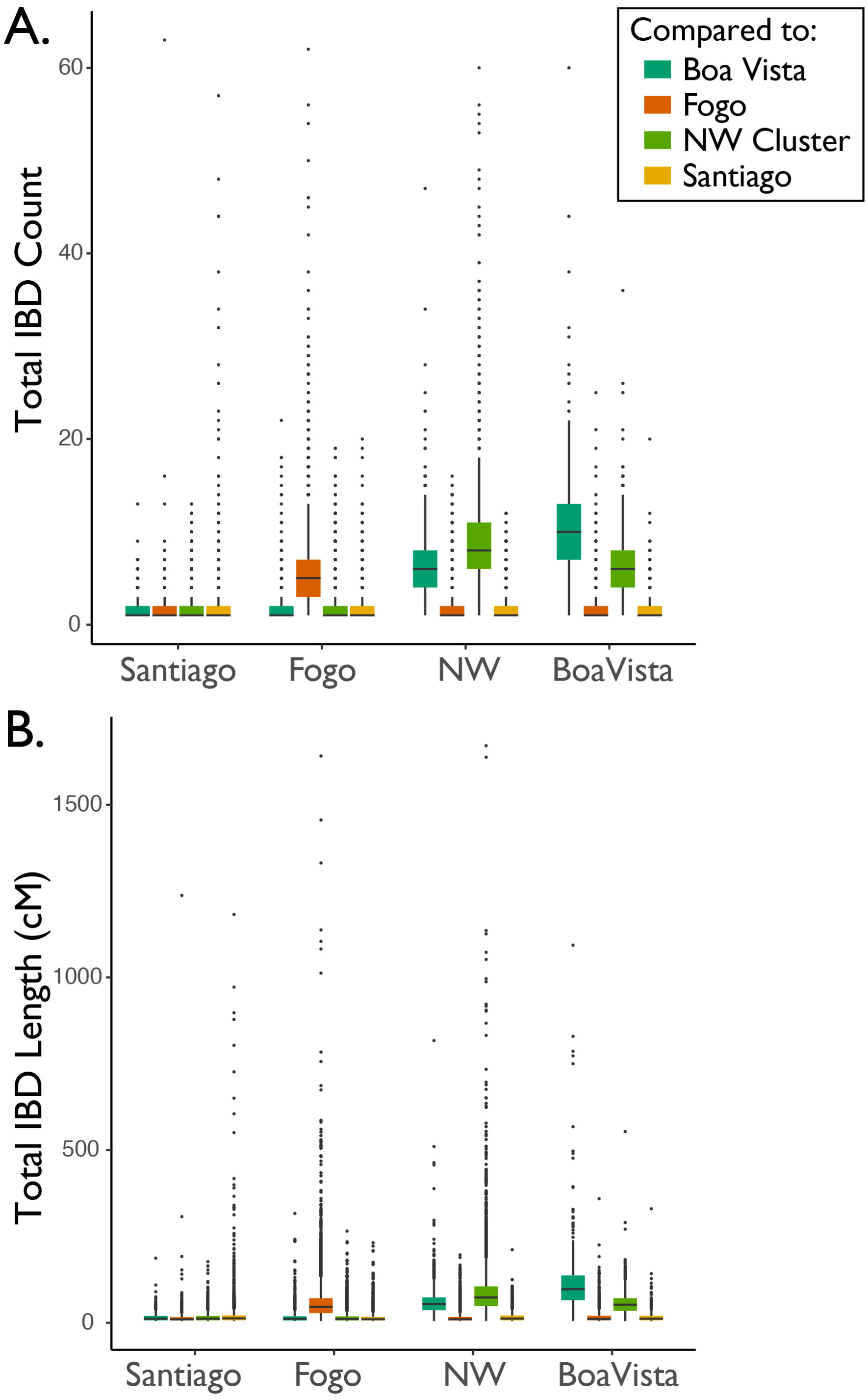


**Supp Fig 5: The distribution of IBD sharing within and between islands.** The distributions of total count (A) and summed length (B) of pairwise IBD segments shared between individuals both within the same island and across islands.

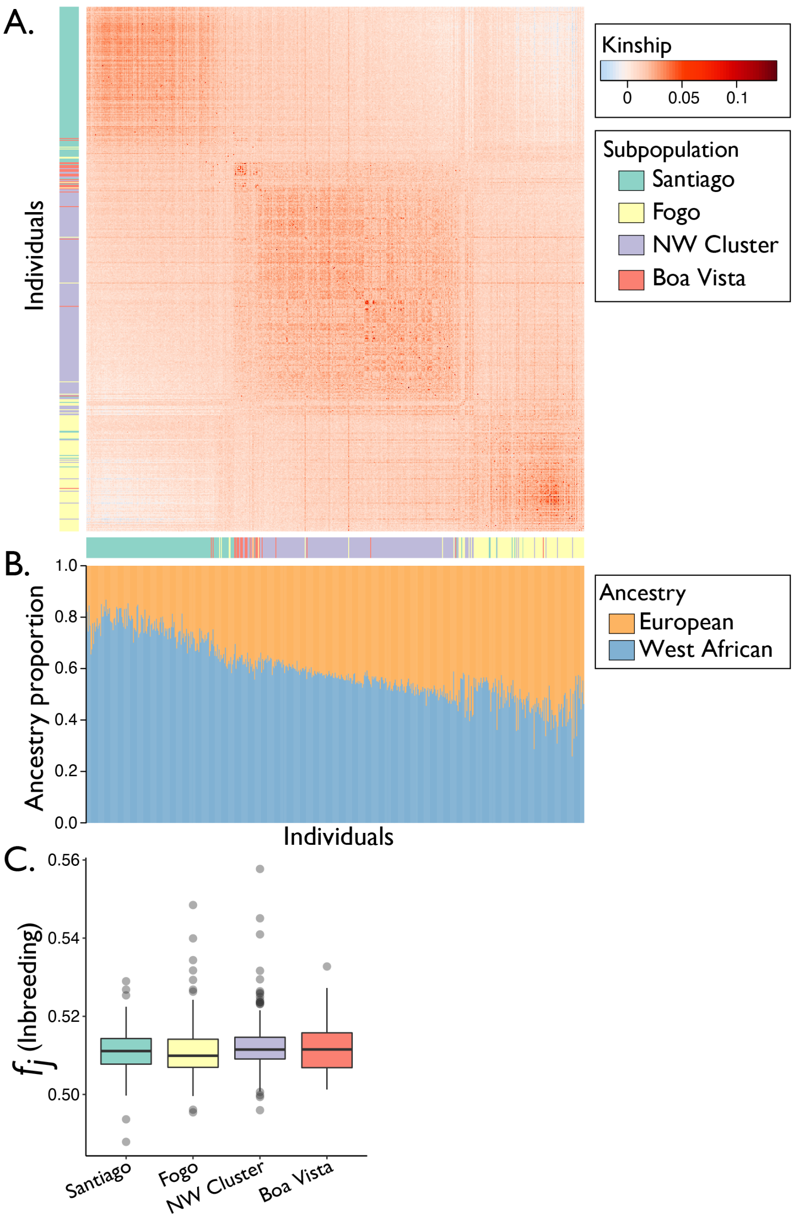


**Supp Fig 6: Kinship estimation in the context of ancestry**. Heatmap (A) showing the full kinship matrix and global ancestry estimates from ADMIXTURE (B) with individuals aligned with the kinship matrix, produced using the method of Ochoa & Storey (2019). Individuals within the matrix were ordered using seriation, which places low kinship values away from the diagonal. The diagonal of the kinship matrix gives the distribution of inbreeding coefficients (C), or $f_j$ as described in Ochoa & Storey (2019), within each island region.

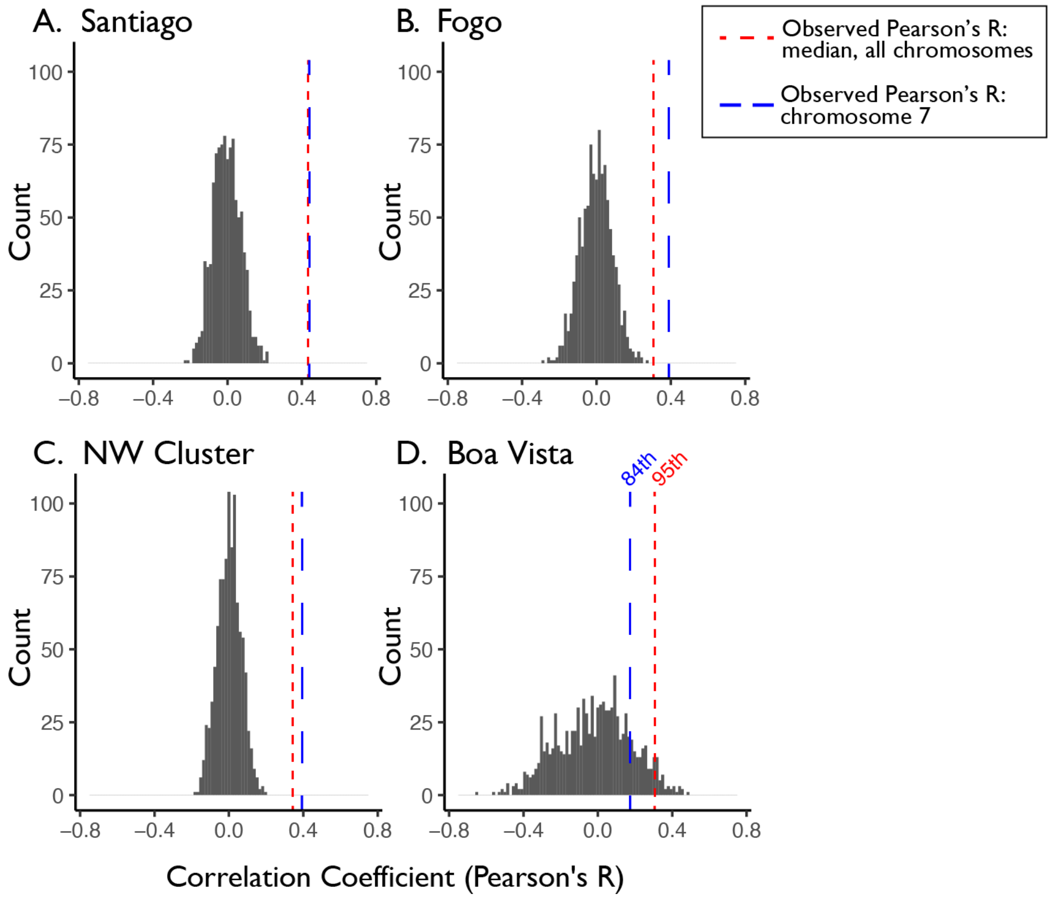


**Supp Fig 7: Comparison of inferred ancestry-assortative mating strength to the sampling distribution based on randomly paired haplotypes.** We compared the empirically-inferred strength of ancestry-assortative mating (red lines above: using the median Pearson's R from the set of all chromosomes shown in Fig 2B) to the correlation observed in random samples. Each distribution includes 1,000 sets of mating pairs that were randomly sampled with replacement from the full set of observed haplotypes in the current generation (using a representative chromosome, chromosome 7). The randomly sampled sets contained the same number of mating pairs as the empirical sample for each island (i.e., 172 pairs per sample for Santiago, 129 for Fogo, 236 for the NW Cluster, and 26 for Boa Vista). For all islands, the empirically-inferred strength of assortative mating differs significantly from the distribution of random samples (Santiago t-test: $t = 176.18$, $df = 998$, $p < 1 \times 10^{-8}$; Fogo t-test: $t = 113.75$, $df = 998$, $p < 1 \times 10^{-8}$; NW Cluster t-test: $t = 171.5$, $df = 998$, $p < 1 \times 10^{-8}$; Boa Vista t-test: $t = 52.07$, $df = 998$, $p < 1 \times 10^{-8}$; significance testing performed using the distribution of differences between empirical Pearson's R (red lines) and Pearson's R in each random sample to test whether the mean of that distribution differs from zero). The empirically-inferred strength of ancestry-assortative mating from a single example chromosome (chromosome 7) is also shown for comparison (blue lines).

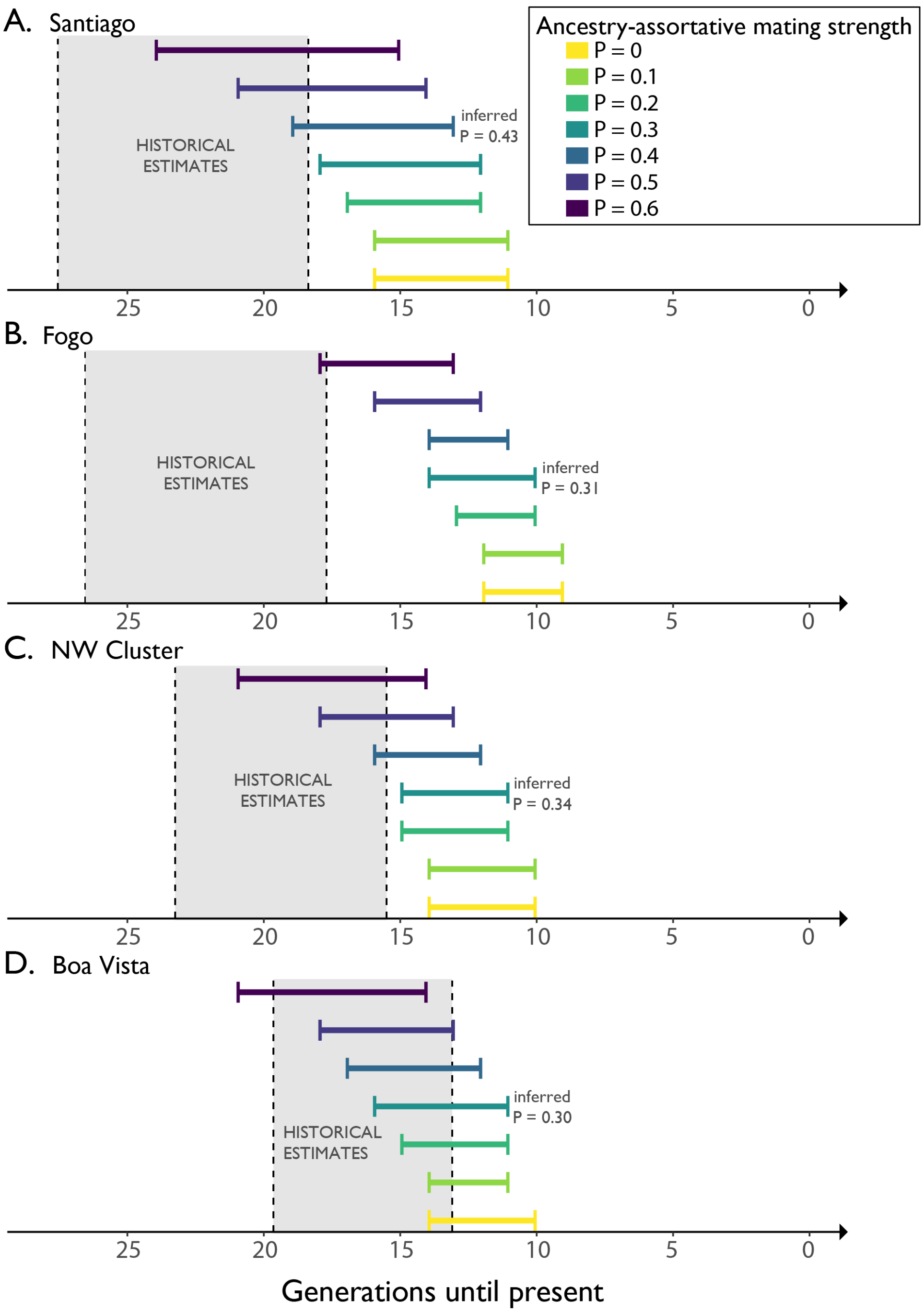


**Supp Fig 8: Timelines of inferred generations of admixture using LAD under varying degrees of ancestry-assortative mating.** For each island, admixture timing estimates inferred with the LAD-based method of Zaitlen et al. (2017) under a range of assortative mating strengths (P = the correlation in ancestries of individuals in mating pairs) are shown in comparison to historical records. Each result includes the range of estimates generated under the assumption of constant migration rate (m = 0.01; yielding older estimates) to the assumption of no migration (m = 0; yielding more recent estimates). For each island, the inferred value of P from Fig 2 is provided as an annotation to the right of the timing estimate that most closely corresponds to the inferred value.

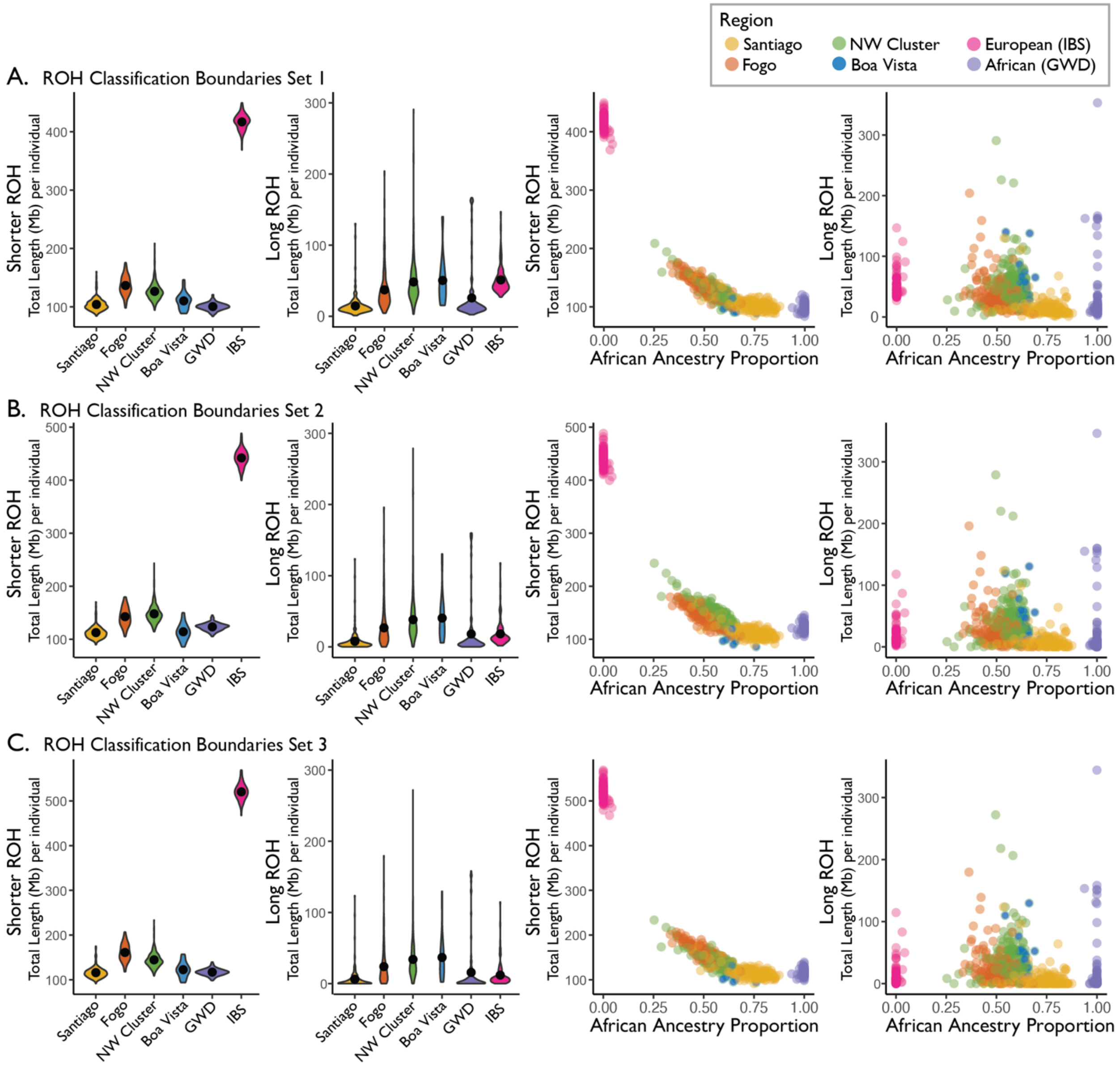


**Supp Fig 9: ROH distributions under various length classification cutoffs.** Repeating Fig 4 under three different sets of ROH length classification rules, the violin plots show the population-specific distributions of the total (summed over each genome) length of autosomal ROH per individual. The scatter plots show the total length of autosomal ROH per individual plotted against West African ancestry proportions and colored by population. (A) Set 1 uses the minimum shorter/long boundary (896,699 bp) reported in Pemberton et al. (2012), (B) Set 2 uses the mean shorter/long boundary (1,548,382 bp) reported in Pemberton et al. (2012), and (C) Set 3 uses the maximum shorter/long boundary (2,191,781 bp) reported in Pemberton et al. (2012).

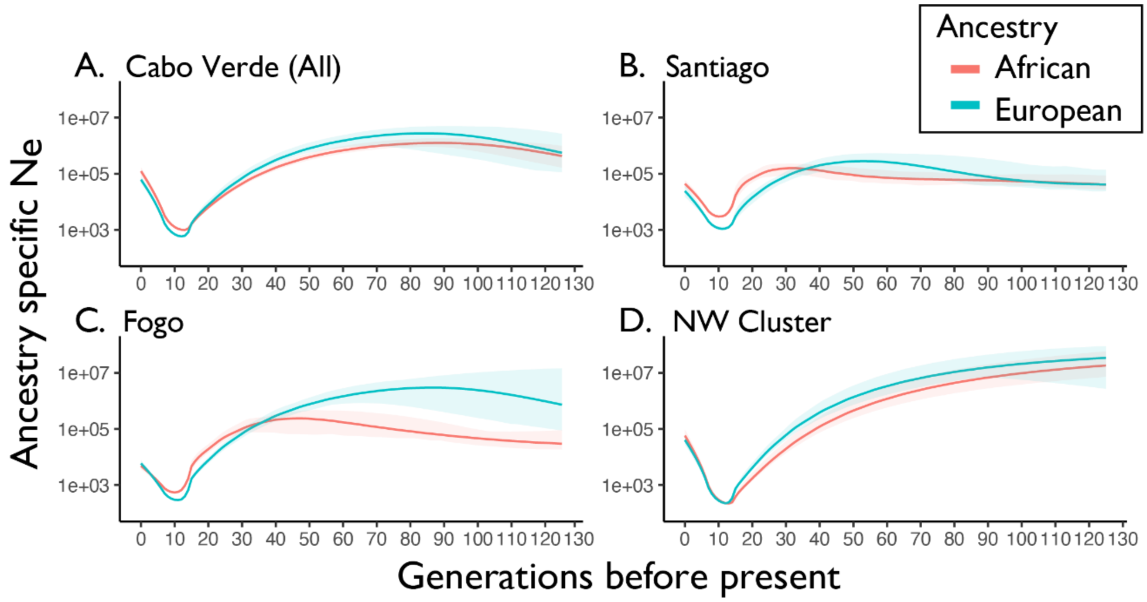


**Supp Fig 10: Ancestry-specific population size**. The estimated effective population sizes (Ne, plotted on a log scale) of West African and European ancestry plotted over time (generations until present). The solid lines show estimated ancestry-specific effective population sizes, and the shaded regions around the lines show 95% bootstrap confidence intervals.

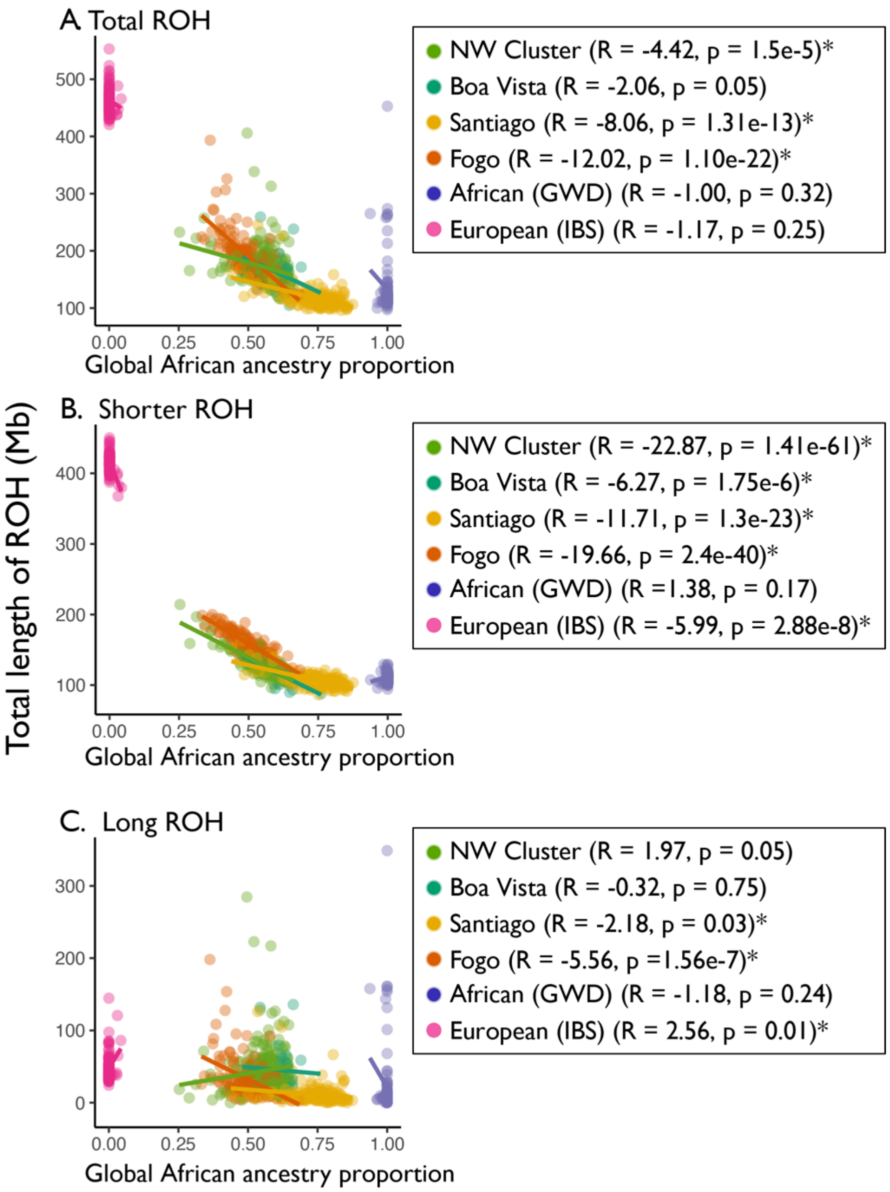


**Supp Fig 11: ROH vs African ancestry proportion**. For all ROH length classes (A), shorter ROH (B), and long ROH (C), the total length of autosomal ROH per individual is plotted against West African ancestry proportions and colored by population. For each length class, ROH within each population is regressed onto global African ancestry proportion.

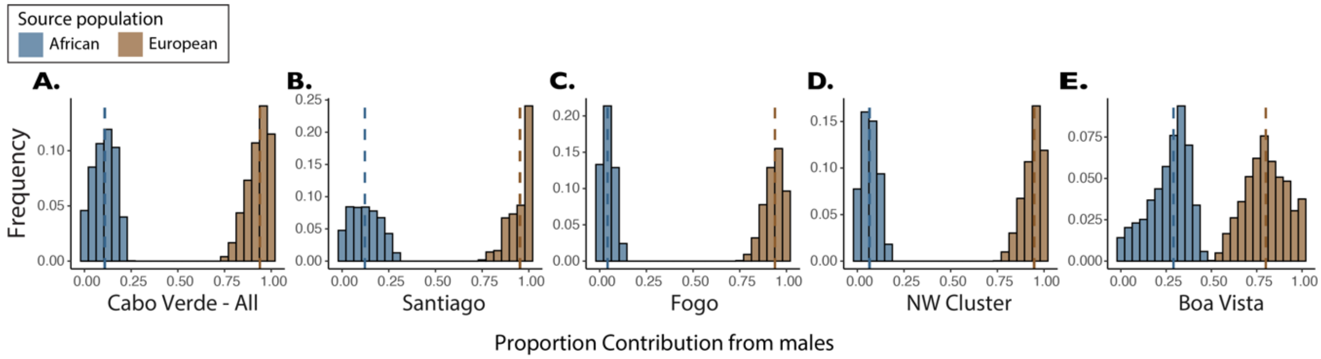


**Supp Fig 12: Sex-biased admixture in Cabo Verde – male contributions.** Under a model of constant admixture over time, the fraction of the total contribution of genetic material originating from males for West African and European source populations. Here, we show the distribution of parameter sets for the smallest 0.1% of Euclidean distances between the model-predicted and observed X and autosomal ancestry from a grid of possible parameter values. The range of sex-specific contributions from West African and European source populations that produce ancestry estimates closest to those observed in Cabo Verde are shown for Cabo Verde as a whole (A), and then broken down by region (B-E), with medians (dashed lines).

**Supp Table 1: Summary of computational methods.**

| Program | Usage in this study | Reference |
|---|---|---|
| *Characterization of ancestry* | | |
| PLINK v1.9 | LD pruning; PCA | Purcell et al. 2007 |
| ADMIXTURE v1.3.0 | Unsupervised estimation of genomic ancestries | Alexander et al. 2009 |
| SHAPEIT v2 | Phasing | Delaneau et al. 2013 |
| RFMix v1.5.4 | Local ancestry calling | Maples et al. 2013 |
| *Inference of admixture timing* | | |
| ALDER v1.03 | Estimation of admixture timing based on the extent of LD decay among neighboring loci | Loh et al. 2013 |
| MULTIWAVER v2.0 | Estimation of admixture timing based on ancestry tracts inferred by RFMix | Ni et al. 2019 |
| LAD-based method | Estimation of admixture timing based on local ancestry disequilibrium (LAD) | Zaitlen et al. 2017 |
| *Testing for assortative mating and sex-biased admixture* | | |
| ANCESTOR | Estimation of the ancestry proportions of the two parents of each individual | Zou et al. 2015 |
| Mechanistic model of sex-biased admixture | Inference of admixture parameters under a model allowing sex-biased contributions from source populations | Goldberg et al. 2015 |
| *IBD, ROH, and relatedness analyses* | | |
| Ancestry specific IBD Ne (ibdne.23Apr20.ae9.jar) | Estimation of ancestry-specific population sizes | Browning & Browning 2018 |
| RefinedIBD (refined-ibd.17Jan20.102.jar) | Inference of segments of IBD | Browning & Browning 2015 |
| Popkin | Kinship estimation under a framework designed for arbitrary population structure | Ochoa & Storey 2019 |
| GARLIC v1.1.6 | Classification of ROH | Szpiech et al. 2017 |

**Supp Table 2: ROH length classification and LOD score cutoffs from Garlic.**

| Population | Class A/B Length Boundary | Class B/C Length Boundary | Window Size | LOD Score Cutoff |
|---|---|---|---|---|
| Santiago | 303,824 bp | 1,081,400 bp | 50 | 2.203 |
| NW Cluster | 308,994 bp | 1,076,210 bp | 50 | 1.746 |
| Fogo | 309,213 bp | 1,118,290 bp | 50 | 0 |
| Boa Vista | 328,023 bp | 1,144,080 bp | 50 | 1.644 |
| African (GWD) | 327,265 bp | 1,262,910 bp | 50 | 1.773 |
| European (IBS) | 264,625 bp | 914,139 bp | 40 | -2.169 |

**Supp Table 3: Estimates of admixture timing.**

| Population | Recorded generations[1] | Inferred generations: ALDER | Inferred generations: MultiWaver | Inferred generations: LAD, random mating | Inferred generations: LAD, assortative mating |
|---|---|---|---|---|---|
| Santiago | 18.37 – 27.55 | 8.81 - 10.05 | 6 – 9; 13 – 15 | 11 – 16 | 13 – 20 |
| NW Cluster | 15.5 – 23.25 | 10.33 – 11.53 | 7 – 10; 13 – 14 | 10 – 14 | 11 – 16 |
| Fogo | 17.7 – 26.55 | 9.74 – 10.52 | 11.5 – 12.5 | 9 – 12 | 10 – 13 |
| Boa Vista | 13.1 – 19.65 | 8.16 – 10.04 | 12 - 13 | 10 - 14 | 11 – 16 |

[1] See *Methods: Historical records* for the sources of historical estimates of admixture timing.
